# Challenging the point neuron dogma: FS basket cells as 2-stage nonlinear integrators

**DOI:** 10.1101/251314

**Authors:** Alexandra Tzilivaki, George Kastellakis, Panayiota Poirazi

**Affiliations:** Institute of Molecular Biology and Biotechnology (IMBB), Foundation for Research and Technology Hellas (FORTH), Heraklion, Greece; Department of Biology, University of Crete, Heraklion, Greece.

## Abstract

Interneurons are critical for the proper functioning of neural circuits. While often morphologically complex, their dendrites have been ignored for decades, treating them as linear point neurons. Exciting new findings reveal complex, non-linear dendritic computations that call for a new theory of interneuron arithmetic. Using detailed biophysical models, we predict that dendrites of FS basket cells in both hippocampus and prefrontal cortex come in two flavors: supralinear, supporting local sodium spikes within large-volume branches and sublinear, in small-volume branches. Synaptic activation of varying sets of these dendrites leads to somatic firing variability that cannot be explained by the point neuron reduction. Instead, a 2-stage Artificial Neural Network (ANN), with sub- and supralinear hidden nodes, captures most of the variance. Reduced neuronal circuit modeling suggest that this bi-modal, 2-stage integration in FS basket cells confers substantial resource savings in memory encoding as well as the linking of memories across time.

---

GABAergic interneurons play a key role in modulating neuronal activity and transmission in multiple brain regions^1–5^. Among others, they are responsible for controlling the excitability of excitatory and inhibitory cells, modulating synaptic plasticity and coordinating synchrony during neuronal oscillations^2,6–8,9^. GABAergic interneurons come in a variety of molecular profiles, anatomical features and electrophysiological properties^1,3,5,10^. Despite this variability, many interneuron types exhibit similar computations, the most common being a precise EPSP-spike coupling^2,11,12^. As they innervate a large number of cells, near the site of action potential initiation, they are believed to generate a powerful widespread inhibition, also referred to as an inhibitory blanket^13^.

Fast Spiking basket cells (FS BCs) constitute one of the main types of hippocampal and neocortical interneurons^6,13,14^. They are part of the PV positive interneuron class, which also includes the axo-axonic, chandelier and bistratified sub-types. FS BCs are distinguished from other subtypes by their anatomical features^15^, synaptic connectivity patterns^13,16^ and membrane mechanisms. These include the presence of calcium permeable AMPA (cp-AMPA) receptors^17,6,18,19^, the low expression of NMDA receptors^19,20^, a weak backpropagation of APs^6,21^, a low density of sodium channels^6^ and a high density of potassium channels in their aspiny dendritic trees^5,15,23,6,4,24^.

A growing body of literature recognizes the importance of FS BCs in controlling executive functions such as working memory and attention as well as their role in neurodegenerative disorders^4,25,26^. However, little is known about the mechanistic underpinnings of FS BC contributions to these functions. Most studies have focused on the molecular and anatomical features of FS BCs^7,13^ and led to the dogma that FS BCs serve as “on-off” cells, integrating inputs like linear –or at best sublinear-point neurons^28,29^.

This dogma is based on the assumption that FS BCs integrate synaptic inputs in a linear manner, completely ignoring potential dendritic infuences^6^. Dendritic integrative properties however, can play a pivotal role in translating incoming information into output signals^30,31,32^. In pyramidal neurons for example, this is often done in highly nonlinear ways that facilitate memory and other executive functions^31,33–36^.

Exciting new findings suggest a potentially similar contribution of dendrites in interneuron function. Sublinear dendritic EPSP integration along with supralinear calcium accumulations has been reported in cerebellar Stellate Cells^11,37^. Moreover, certain interneuron sub-types in the CA1 area exhibit dendritic supralinearities^38,39^ while in the CA3, both calcium nonlinearities and sodium spikes in FS BC dendrites during sharp wave ripples have been reported^2^. The exact nature of dendritic computations in FS BCs, however, is unknown. As a result, whether a linear point neuron or a more sophisticated abstraction -like the two-stage^40^ or multi-stage integration proposed for pyramidal neurons-can successfully capture their synaptic integration profile, remains an open question.

To address these questions, we developed an elaborate toolset that consists of a) detailed, biologically constrained biophysical models of hippocampal and cortical FS BCs, b) reduced 2-stage integrate-and-fire models of these cells, c) 2-layer Artificial neural network abstractions and d) a large scale microcircuit model of 2-stage pyramidal, FS BC and dendrite targeting (SOM) interneurons (See **Online Methods** and **Figure 1**). We first characterized the integration profiles of FS BC dendrites using the detailed biophysical models. Synaptic stimulation predicted the co-existence of two distinct modes within the same tree: some dendrites exhibited supralinear while others sublinear summation of inputs (**Figure 2, Supplementary Figures 4, 5**). Supralinear dendrites supported local, sodium-dependent spikes (**Supplementary Figure 7**) and were characterized by large volume and low input resistance (**Figure 3**), which are shaped by the combination of dendritic length and diameter. Direct manipulation of these anatomical features in biophysical models gated the induction of sodium spikes and determined the integration mode (**Figure 3**). Using an array of different activation patterns, we found that spatially dispersed inputs lead to higher firing rates than inputs clustered within a few dendrites (**Figure 4**), opposite to respective findings in pyramidal neurons^33^. Moreover, these different activation patterns result in a wide range of firing rates that are better explained by a 2-layer Artificial Neural Network (ANN) with non-linear hidden layer activation functions rather than a linear ANN (**Figures 5, 6, Table 1**). Finally, in order to assess the functional implications of these predictions, we built a reduced network model^41^ of 2-stage integrator neurons (**Figure 1D**) and showed that bi-modal nonlinear integration in FS BCs is beneficial for memory engram storage as well as the linking of memories across time (**Figure 7**).

**Table 1.** ANN regression performance (R^2^) for individual sets of synapses in the 8 model cells. Comparison of ANN prediction accuracy (measured as the R^2^) for linear and 2-layer modular ANN reductions across all 8 FS BC models, tested on three sets of synaptic inputs consisting of 20, 40 or 60 activated synapses, respectively. Synapses were randomly distributed in various ways/locations in the biophysical model cells and resulting firing rates were used as target vectors for the ANNs. The 2-layer modular ANN is clearly superior to the Linear ANN when it comes to capturing location-induced firing-rate variability.

| **Table 1. ANN regression performance ($R^2$) for individual sets of synapses in the 8 model cells** |  |  |  |
| --- | --- | --- | --- |
| ANN type | 20 synapses | 40 synapses | 60 synapses |
| 2-layer modular ANN | 0.8224 | 0.7432 | 0.6167 |
|  | 0.8228 | 0.9060 | 0.8339 |
|  | 0.8048 | 0.8709 | 0.7971 |
|  | 0.7951 | 0.8563 | 0.8127 |
|  | 0.8242 | 0.8352 | 0.7752 |
|  | 0.9098 | 0.8857 | 0.8921 |
|  | 0.8879 | 0.8975 | 0.8827 |
|  | 0.8415 | 0.8600 | 0.7801 |
| Linear ANN | 0.4541 | 0.3484 | 0.2856 |
|  | 0.6814 | 0.7462 | 0.5966 |
|  | 0.6436 | 0.5201 | 0.5625 |
|  | 0.5636 | 0.4475 | 0.4768 |
|  | 0.5550 | 0.4919 | 0.4707 |
|  | 0.7832 | 0.7242 | 0.6758 |
|  | 0.6263 | 0.5463 | 0.4400 |
|  | 0.6179 | 0.7039 | 0.4991 |

**Figure 1.**
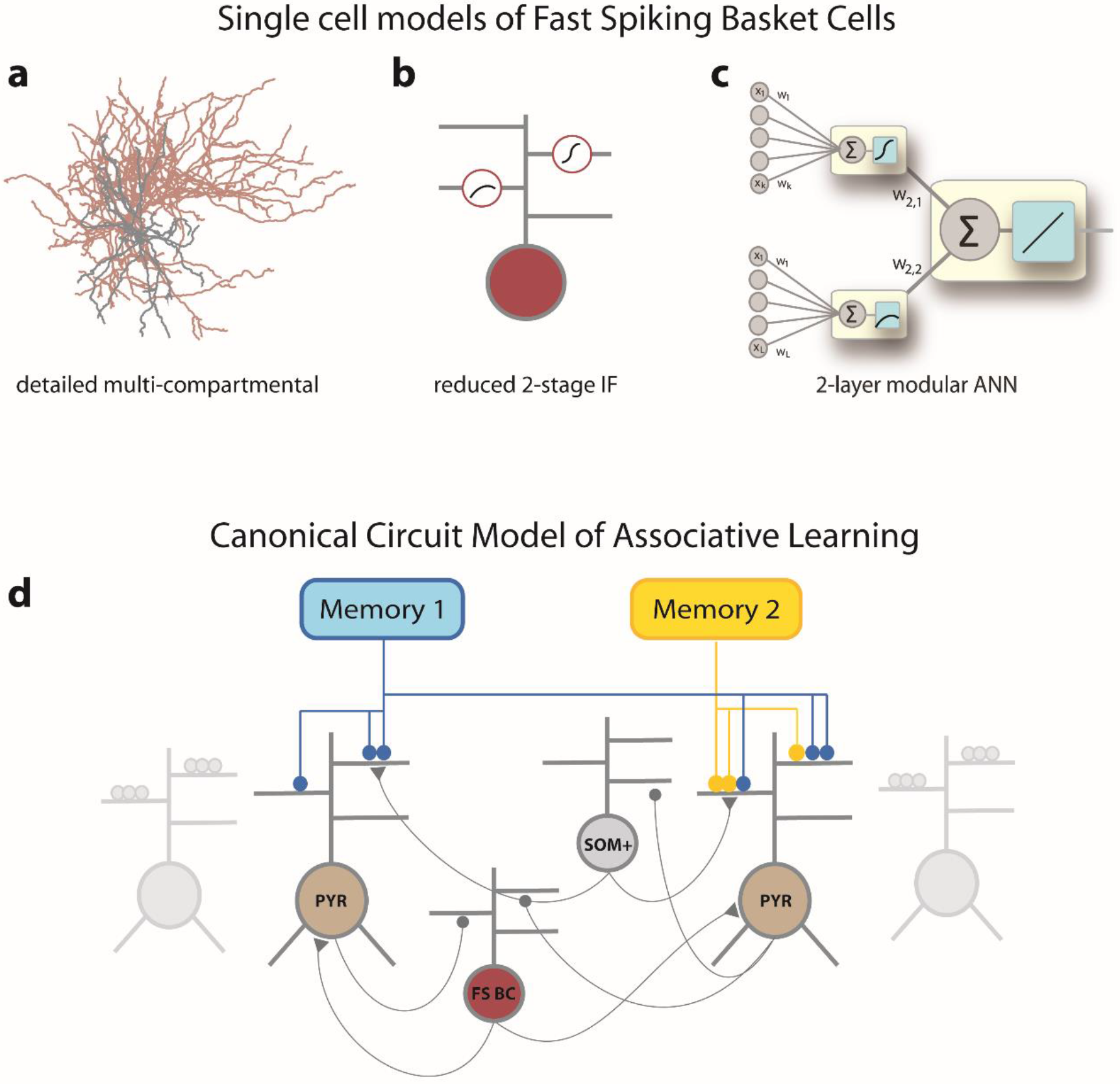
Modeling tools used to study dendritic integration in FS BCs and its functional implications. **a**) Detailed, biophysically constrained multi-compartmental models using realistic reconstructions. **b**) Reduced 2-stage integrate and fire models of FS BCs. **c)**2-layer ANN reduction describing the FS BCs **d**) Reduced network model with simplified pyramidal, FS BCs and SOM+ interneurons. FS-BCs and SOM+ interneurons provide feedback inhibition to excitatory neurons (beige), with FS-BCs interneurons targeting the somatic subunit while SOM+ neurons target the dendritic subunits. Memory encoding afferents provide inputs to excitatory cell dendrites.

**Figure 2.**
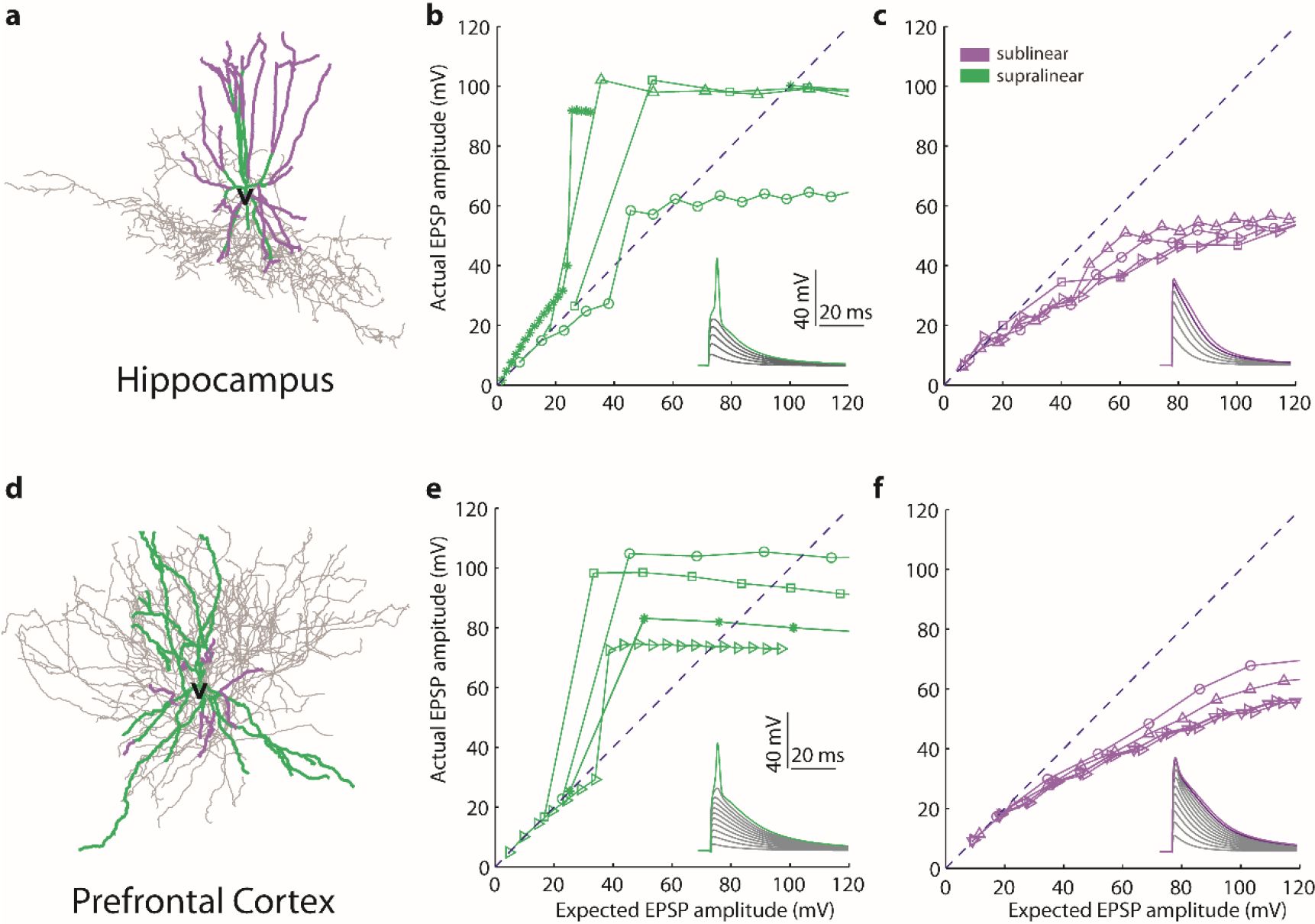
Bimodal dendritic integration in multi-compartmental FS BC models. Examples of Hippocampal (**a**) and PFC (**d**) FS BC morphological reconstructions. Representative input-output curves from supralinear (**b**, **e**) and sublinear (**c**, **f**) dendritic branches in Hippocampal (top) and PFC (bottom) models, in response to synaptic stimulation. Increasing numbers of synapses (from 1 to 20 with step=1) are uniformly distributed within each stimulated branch and are activated with a single pulse. The y-axis shows the amplitude of the dendritic EPSP caused by synaptic activation while the x-axis shows the expected EPSP amplitude that would result from the linear summation of synaptic EPSPs. The dashed line indicates linear summation. Insets show representative traces.

**Figure 3.**
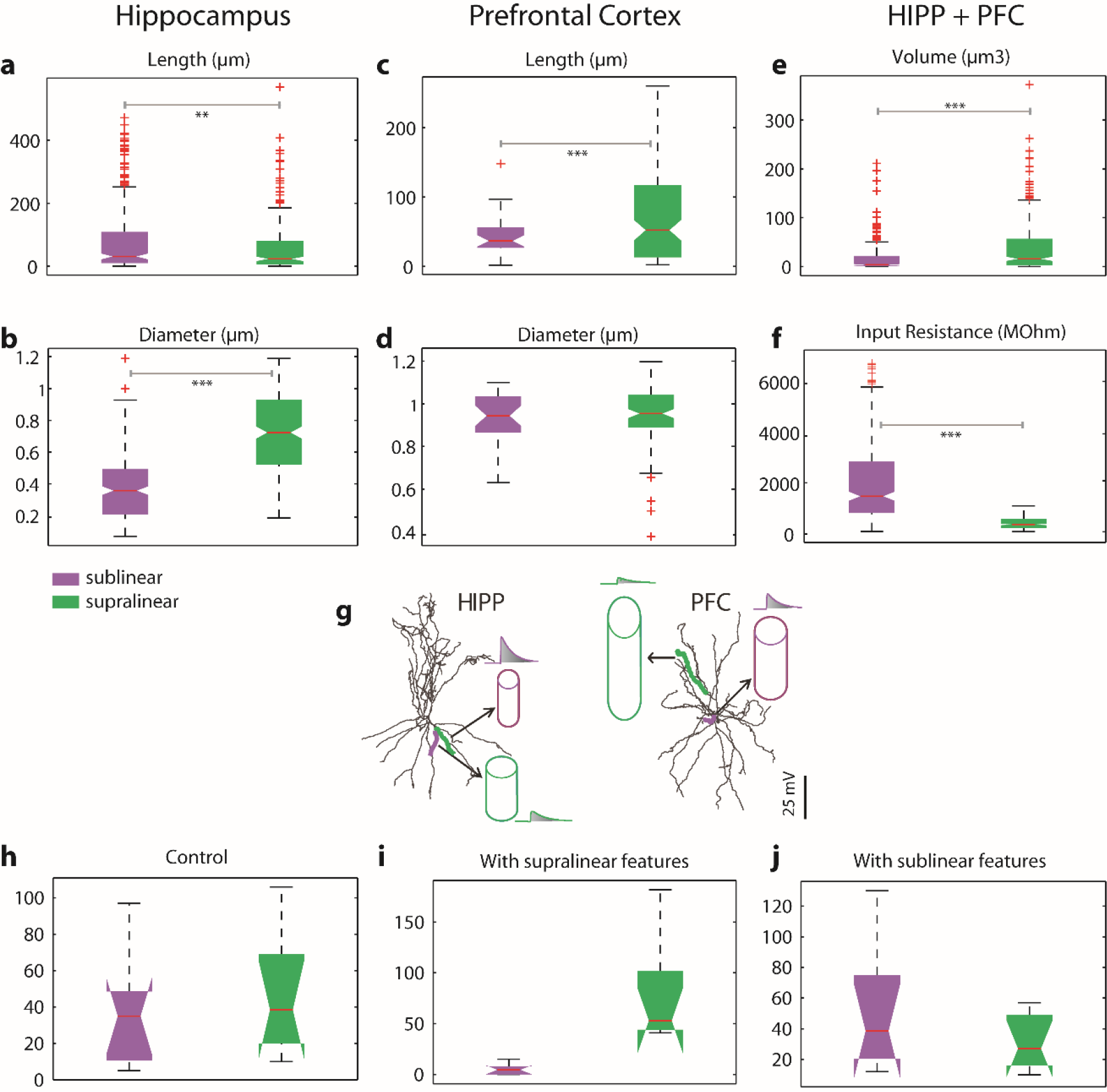
Morphological determinants of dendritic integration mode. **a**, **c**: Total length distributions of supralinear vs. sublinear dendrites in the hippocampus (**a**) and the PFC (**c**). Statistically significant differences are observed for both sub- and supra-linear dendrites, in both areas. (p-value<0.0001 for Hippocampus and p-value<0.01 for PFC). **b**, **d**: Same as in **a**, **b**, for mean dendritic diameter. Statistically significant differences are observed in Hippocampal (p-value<0.0001) but not in PFC FS BCs. **e**-**f**. Dendritic Volume and dendritic Input Resistance are common discriminating characteristics among supralinear (larger, with low input resistance) and sublinear (smaller, with high input resistance) dendrites, for both areas (p-value<0.0001 for Hippocampus and PFC, for volume and Input Resistance respectively). **g**. Schematic illustration of morphological features for supralinear and sublinear dendrites in Hippocampus (left) and PFC (right). Traces indicate first EPSP in supralinear and sublinear dendrites. **h**-**j**. Distributions of the number of supralinear and sublinear dendrites in both areas, under control conditions (**h**), with the mean diameter and length of all dendrites set to the mean values of the supralinear class (**i**) and with mean diameter and length of all dendrites set to the mean values of the sublinear class (**j**).

**Figure 4.**
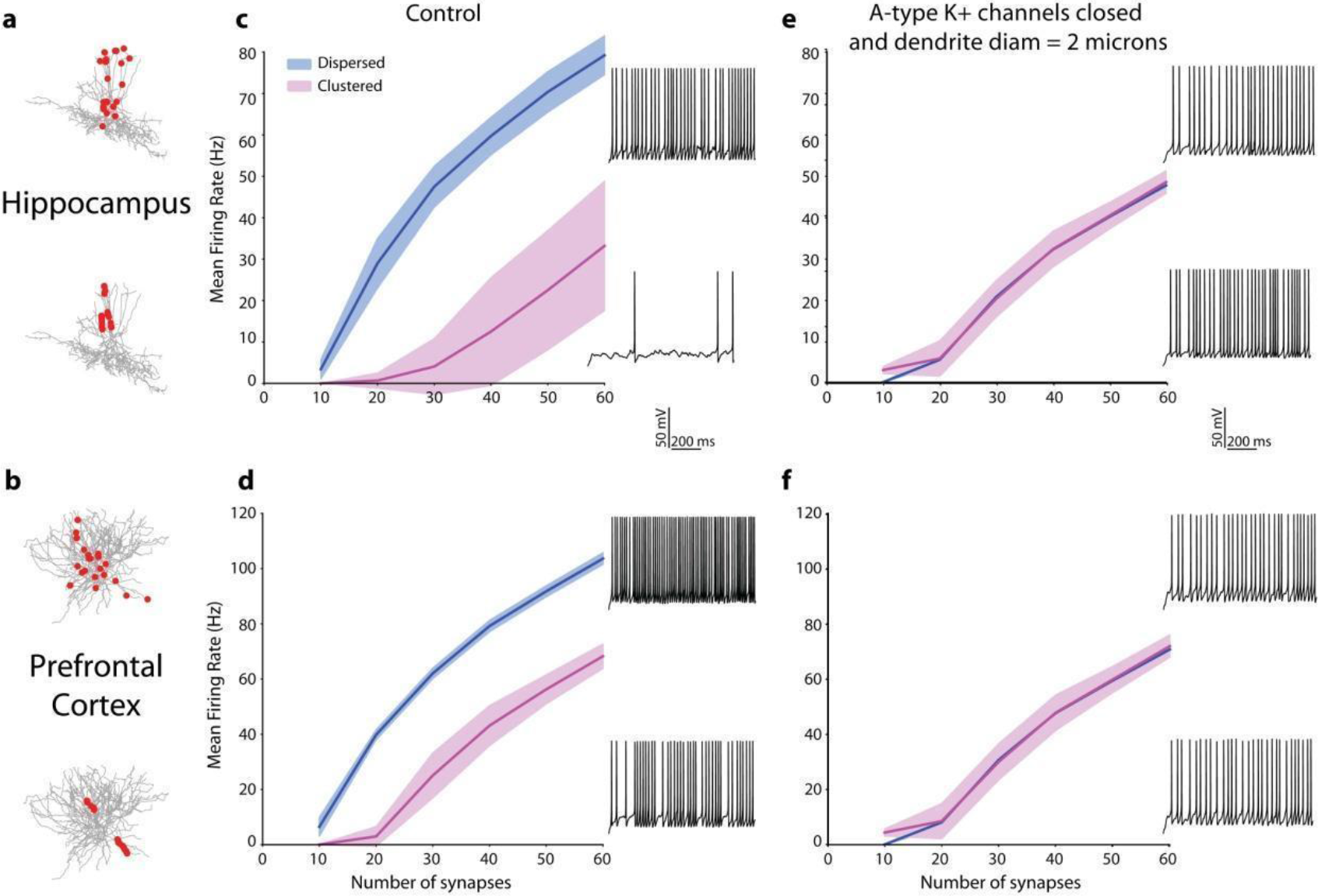
Effect of bimodal dendritic integration on neuronal firing. Firing rate responses (in Hz) from one Hippocampal (**a**,**c**) and one PFC (**b**,**d**) model cell, in response to stimulation of increasing numbers of synapses (10 to 60) that are either randomly distributed throughout the entire dendritic tree (blue) or clustered within a few dendritic branches (pink).) Synapses are stimulated with a 50 Hz Poisson spike train. In both cases, dispersed activation leads to higher firing rates. **e**,**f**: Same as in **c**,**d**with dendritic diameter set to 2 microns and removal of A-type dendritic channels. Firing rates are indistinguishable between clustered and dispersed activation patterns. Insets depict representative traces from dispersed (top) and clustered (bottom) activation of 30 synapses 30. Red dots in show the synaptic allocation motif in **a**, **b**.

**Figure 5.**
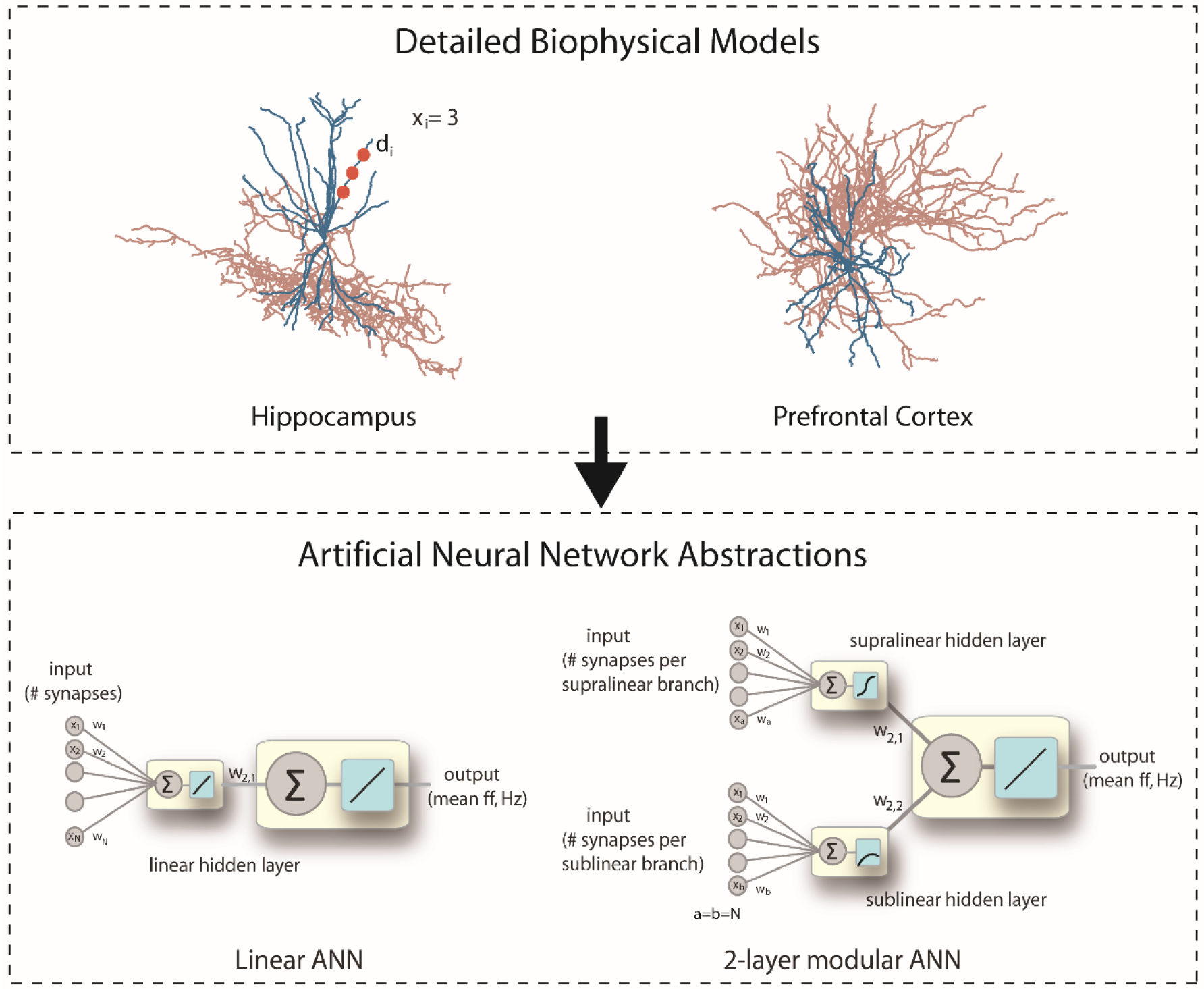
Reduction of multi-compartmental models into ANN abstractions. Two types of abstractions are examined: **a**) a Linear ANN, in which the input from all dendrites (xi=number of synapses in dendrite i, N=number of dendrites) is linearly combined at the cell body and **b**) a 2-layer modular ANN, in which the input is fed into two parallel, separated hidden layers. The supralinear-layer receives the number of inputs landing onto supralinear branches (a=number of supralinear dendrites) while the sublinear layer receives the number of inputs landing onto sublinear dendrites (b=number of sublinear dendrites). Neurons in both hidden layers are equipped with nonlinear transfer functions, a logistic sigmoid in the supralinear layer and a sublinear function in the sublinear layer. The somatic transfer functions of both ANNs are linear.

**Figure 6.**
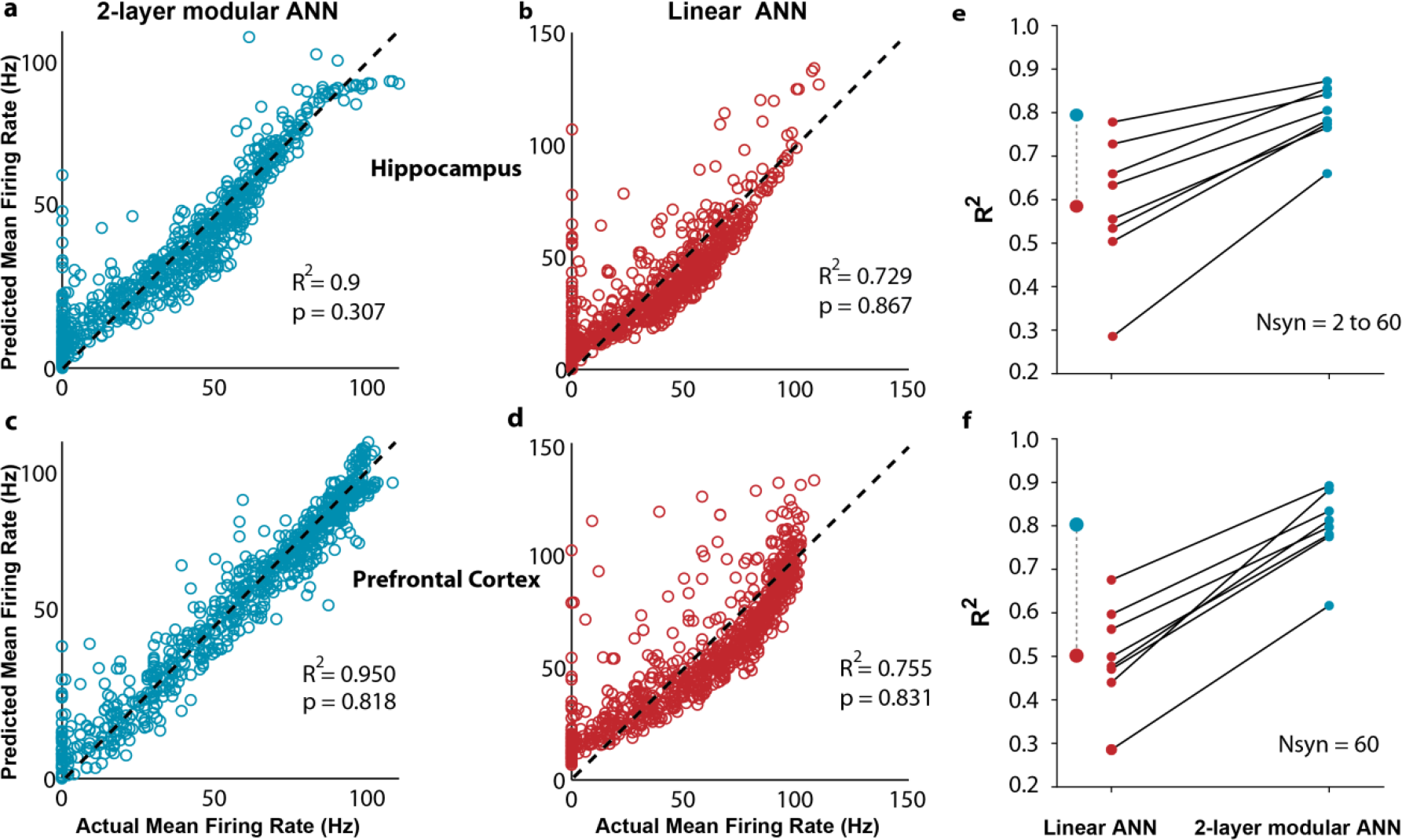
Challenging the point neuron dogma: FS basket cells as 2-stage nonlinear integrators. Linear regression analysis for 2-layer modular (**a, c**) and linear (**b, d**) ANNs for one indicative Hippocampal (top) and one indicative PFC (bottom) model cell. Actual Mean Firing Rates (Hz) correspond to the responses of the compartmental model when stimulating with 50Hz Poisson spike trains-varying numbers of synapses (1 to 60), distributed in several ways (clustered or dispersed) within both sub- and supra-linear dendrites. Expected Mean Firing Rates (Hz) are those produced by the respective ANN abstraction when receiving the same input (number of stimulated synapses) in its respective sub-/supra- or linear input layer nodes. **e)**Regression performance (measured as R^2^) for 2-layer modular (right) and Linear (left) ANNs for all 8 FS BC model cells respectively. In all cases the 2-layer modular ANNs is superior to the Linear ANNs. Mean R^2^ values over all cells for the Linear (red) and 2-layer modular (cyan) ANNs are shown in the right. f) Same as e), applied to datasets comprised of 60 input synapses. The difference in performance of the two ANN types is higher in this challenging task.

**Figure 7.**
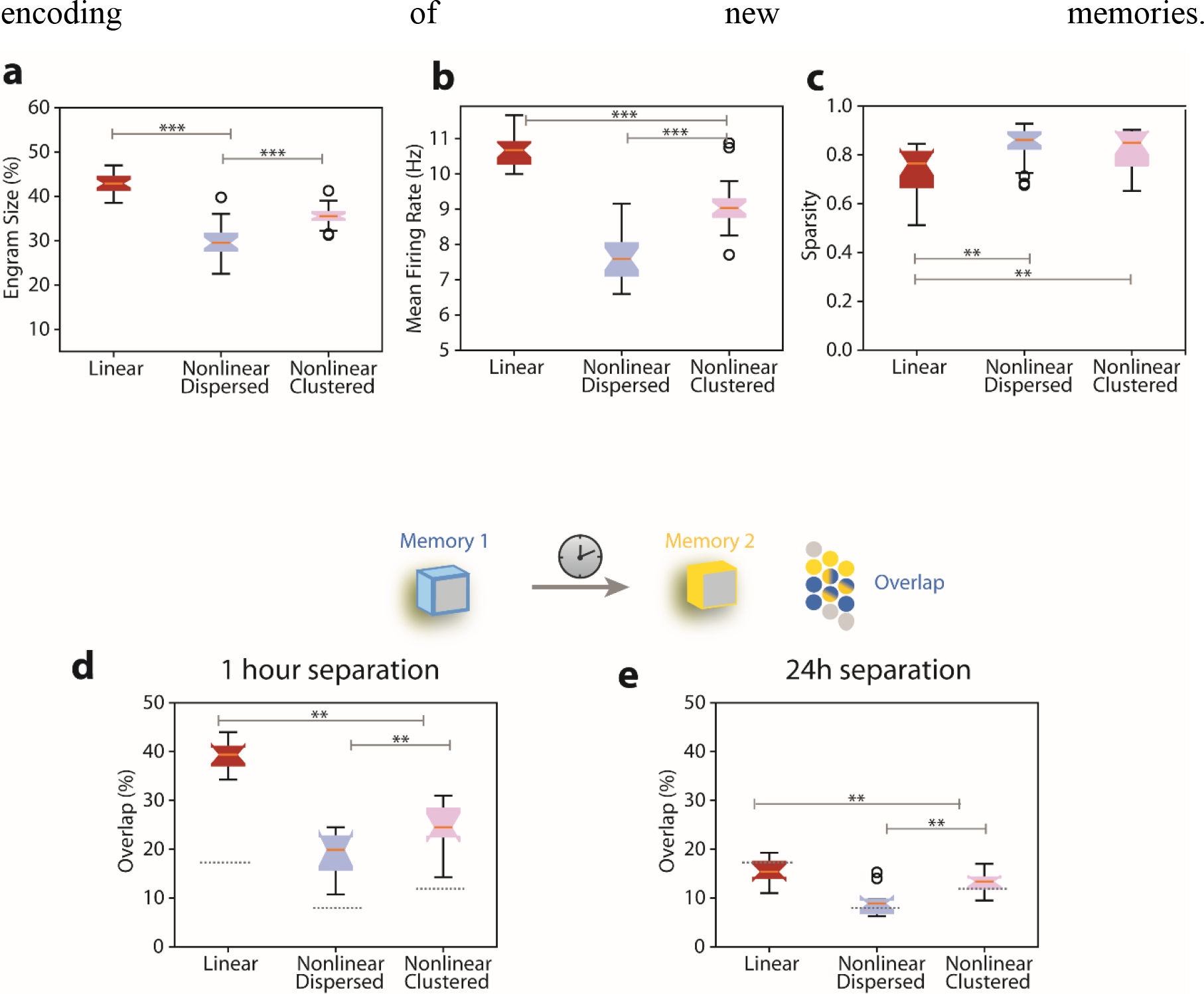
Properties of memory engram encoding under different dendritic nonlinearity configurations, using the circuit model depicted in Figure 1d. **a**) Size of memory engram (percentage of excitatory neurons that respond with *ff*>10Hz during memory recall) for Linear/Bi-modal FS-BC dendritic subunits receiving dispersed (blue) or clustered (pink) synaptic inputs. **b**) Mean firing rate of the excitatory population under the conditions enumerated in (**a**). **c**) Treves-Rolls sparsity metric of the excitatory population firing rates under the conditions enumerated in (**a**). **d**) Percentage of overlap between two memory engrams when 2 memories are separated by 1 hour, under the conditions enumerated in (B). Dashed lines indicate the chance level of overlap for the engram sizes of the dispersed case shown in (**a**). **e**) As in (**d**) for 24 hours separation. Box plots indicate data from 20 simulation trials for A-D, and 10 trials for E-F. **: *p<0.05,* *** *p < 0.005*.

This work provides a systematic, cross-area analysis of dendritic integration in FS BCs and its functional implications. Our findings challenge the current dogma, whereby interneurons are treated as linear summing devices, essentially void of dendrites. We predict that the dendrites of FS BCs in both Hippocampal and Neocortical regions can operate in distinct non-linear modes. As a result, FS BCs, similar to pyramidal neurons^40^, are better represented by a 2-stage integrator abstraction rather than a point neuron. Importantly, non-linear dendritic integration in these cells offers substantial advantages for memory encoding in large scale networks.

## RESULTS

### Multi-compartmental, biophysical models

A total of 8 biophysical model neurons were built using realistic reconstructions of FS BCs from rat hippocampal areas (5 cells) and from the prefrontal cortex of mice (3 cells) (**Supplementary Figure 5**). To ensure biological relevance, ionic and synaptic conductances as well as basic membrane properties of model cells were heavily validated against experimental data^6,13,16,23^ (**Supplementary Table 1-4, Supplementary Figures 1–3**). Moreover, for consistency reasons, the same set of biophysical mechanisms (type and distribution) was used in all model cells.

### Bi-modal dendritic integration in Fast Spiking Basket cells

The first step for deducing a realistic abstraction of FS BCs is the systematic characterization of dendritic/neuronal integration properties across a significant number of neurons and dendrites. Towards this goal, we simulated gradually increasing excitatory synaptic input to the dendrites of all neuronal models and recorded the voltage response both locally and at the soma^11,44^. Increasing numbers of synapses (1-20) were uniformly distributed in each stimulated dendrite and activated synchronously with a single pulse. For this particular experiment, sodium conductances in somatic and axonal compartments were closed to avoid backpropagation contamination effects^55,42^ that were detectable in some dendrites. We compared measured EPSPs to their linearly expected values, given by the number of activated inputs multiplied by the unitary EPSP. We found that within the same dendritic tree, branches summate inputs either in a supralinear or a sublinear mode (**Figure 2, supplementary Figures 4, 5**). While there were differences in the number of dendrites and proportions of sub-vs. supralinear dendrites, all of the morphologies tested expressed both integration modes (**Supplementary Table 5**). Moreover, while both modes have been suggested in distinct interneuron types^11,38^, their co-existence in the same tree has yet to be reported.

To assess the robustness of this finding, we first performed a sensitivity analysis whereby the cp-AMPA, NMDA, VGCCs, sodium and A-type potassium conductances were varied by ±20% of their control value. We found no changes in the integration mode of dendrites (data not shown) and only insignificant alterations in the spike threshold of supralinear dendrites (**Supplementary Figure 6b**). The only manipulation that eliminated supralinearity was the blockade of dendritic sodium channels (**Supplementary Figure 7**).

Next, we examined whether the two modes are influenced by the presence of Gap junctions, which are well established in FS BCs^46^. Towards this goal, we connected pairs of Hippocampal and PFC cells with 10 electrical synapses (see **Online Methods**). Presynaptic cells were synaptically activated so as to fire at gamma rate frequency as per Tamas et al 2000^46^ and the integration mode was assessed, as previously, in the dendrites of the post-synaptic cell. We found no influence of gap junctions on the integration mode, apart from a slightly increased membrane potential (**Supplementary Figure 8**).

The same effect was observed in simulations of more physiological conditions such as active whisking^47^. This was done via weak synaptic activation of randomly selected dendrites resulting in a somatic firing rate of 3±1 Hz^47^ (see **Online Methods**). Both modes of dendritic integration remained unaffected by the presence of *in vivo* like activity fluctuations (**Supplementary Figure 9**).

Taken together, the above simulations establish the robustness of bi-modal dendritic integration and suggest that under physiological conditions, FS BCs are likely to express both types of dendritic integration modes.

### Determinants of dendritic integration modes

Next, we searched for biophysical and/or anatomical determinants of the two integration modes. Blockade of sodium conductances in the dendrites eliminated the supralinear integration mode in all morphologies tested (**Supplementary Figure 7**), but this was not the case for blockade of cp-AMPA, NMDA, VGCCs or A-type potassium channels. (**Supplementary Figure 6a**). These simulations indicate that sodium channels are the key ionic mechanism underlying the supralinear mode. What remains unclear is why these model cells also have sublinear dendrites, when the distribution and conductance values of sodium channels is the same in all dendrites.

Since morphological features of dendrites were previously shown to influence synaptic integration profiles^36^, we investigated whether anatomical features correlate with the expression of each integration mode. We found that the mean dendritic diameter was highly statistically different (p-value=2.6041e-60) among sub-(thinner) and supra-linear (thicker) dendrites in the hippocampus (**Figure 3b**) while in the PFC the dendritic length was a better determinant of sub-(shorter) vs. supra-linearity (longer) (p-value=4.1768e-04) (**Figure 3c**). Length was less, yet, important in the hippocampus (p-value=0.0040) (**Figure 3a**) while diameter was not different among sub- and supralinear dendrites in the PFC (p-value=0.9458) (**Figure 3d**). Dendritic volume and input resistance consider both of the above anatomical features and serve as robust morphological / electrophysiological determinants for all dendrites in both areas (p-value=9.8516e-11, 3.9457e-45 respectively), (**Figure 3e,3f**).

Overall, we found that supralinear dendrites have high volume and low input resistance while sublinear dendrites have smaller volume and high input resistance (**Figure 3e, 3f**). This can be explained by considering the fast kinetics of cp-AMPA receptors and A-type potassium channels in the dendrites of FS BCs. In sublinear dendrites, where the input resistance is high (small volume), coincident synaptic input induces a large, fast rising EPSP which in turn strongly activates the A-type potassium channels that rapidly repolarize the membrane, thus preventing the branch from spiking^12^. The opposite is true for supralinear dendrites, where the low input resistance results is smaller depolarizations that drive smaller A-type potassium currents, enabling the branch to reach the sodium spike threshold. This explanation is consistent with prior findings^2,6,42,12^.

To test the above proposition, we performed causal manipulations whereby we fixed the diameter and length of all dendrites to the mean values of first the supralinear and then the sublinear class and assessed the effect on integration mode. We found that setting the dendritic anatomy to that of a given class also dictated the integration mode (**Figure 3h-j**). These findings suggest that, under the experimentally constrained conductance values for sodium channels, morphology plays a crucial role in the ability of a given dendrite to support local sodium spikes and express the supralinear integration mode.

### Effect of bimodal dendritic integration on neuronal firing

To assess the impact of bi-modal dendritic integration on neuronal output, we simulated a large variety of different spatial patterns of synaptic activation and measured the resulting firing rates. Specifically, we generated over 10,000 synaptic stimulus patterns, which comprised of increasing numbers of excitatory synapses. Synapses were either placed within a few, strongly activated branches (clustered) or they were randomly distributed within the entire dendritic tree (dispersed). In all cases, synapses were activated with random Poisson spike trains at 50 Hz (see **Online Methods**). Dendrites were selected at random and inputs were distributed uniformly within selected dendrites. For the dispersed case, we allocated 2, 5, or 10 synapses in randomly selected dendrites, one at a time, while for the clustered case we allocated 10,15,20,30,60 synapses within an increasing number of branches. In all cases, the number of activated synapses increased gradually up to a maximum of 60, as this number was sufficient to induce spiking at gamma frequencies (30-100 Hz). This process was repeated k times (k = number of dendrites in each cell) to ensure full coverage of the entire tree. As expected given the two modes of dendritic integration, the localization of activated inputs affected neuronal firing. For a given number of activated synapses, dispersed activation led to *higher* somatic firing rates than clustered activation, particularly during gamma related frequencies (30-100 Hz) both in Hippocampal (**Figure 4c**) as well as in PFC FS basket cells (**Figure 4d**). Interestingly, this finding is opposite to what has been reported for pyramidal neurons, in which synapse clustering increases firing rates^33^.

It was previously proposed that the combination of a small diameter with an increased conductance of A-type potassium channels in FS BCs underlies the preference for dispersed synaptic allocationions^6^. To tests this hypothesis, we repeated the above experiment after increasing the diameter (to 2 microns) and blocking the A-type potassium conductance in all dendrites. As shown in **Figure 4e-f**, this manipulation resulted in very similar firing rates irrespectively of the spatial arrangement of synapses, thus eliminating the preference for dispersed allocation of excitatory inputs.

**Supplementary Figure 12** shows the relative contributions of these two mechanisms in our model cells. Disperse synaptic arrangements benefit mostly from the dendritic morphology of FS BCs, as setting the dendritic diameter to 2 microns sharply decreases this preference (**Suppl. Fig. 12 a, b**). This is likely because small diameters prevent signal loss, enabling the small depolarizations produced by dispersed inputs to reach and excite the soma. Clustered arrangements on the other hand, are severely hampered by the high conductance of the A-type potassium channels^24^, as blockade of these currents enhances somatic output (**Suppl. Fig. 12 c, d**). This is because clustered -but not disperse-inputs induce large dendritic depolarizations which strongly activate A-type channels. Since NMDA currents^7^, which would further boost and prolong the cluster-induced EPSPs, are very small in these neurons, the hyperpolarizing effects of the A-type currents are larger than the depolarizing effects of clustered activation.

Another factor that contributes to disperse preference, is dendritic integration. Unlike pyramidal neurons where dendrites are mostly supralinear and benefit from clustered inputs via the induction of dendritic spikes^31,35,40^, these neurons also have sublinear dendrites which dampen the abovementioned benefit. The higher the percentage of sublinear dendrites, the larger the dampening, as: 1) the probability of allocating clustered inputs in the few supralinear dendrites is much smaller and 2) activating sublinear dendrites with clustered inputs offers little/no advantage as dendritic spikes don’t occur in these branches. As shown in **Supplementary Table 7** the more sublinear dendrites a FS BC model has, the weaker the response to clustered input.

Taken together, this analysis reveals that the combination of a high conductance of A-type channels (which penalizes clustering), the specific morphological features of FS BCs (which favor dispersed inputs), and the presence of multiple sublinear dendrites underlie the preference of these cells for disperse rather than clustered activation of their inputs, contrary to pyramidal neurons^40^.

### FS basket cells as 2-layer artificial neural networks

The non-linear synaptic integration taking place within the dendrites of cortical^48^ and CA1^35,44^ pyramidal neurons was previously described as a sigmoidal transfer function^49^. Based on this reduction, a single pyramidal neuron was proposed to integrate its synaptic inputs like a 2-layer artificial neural network, where dendrites provide the hidden layer and the soma/axon the output layer^40^. To assess whether a similar mathematical formalism could be ascribed to FS BC models, we constructed linear and non-linear artificial neural networks (as graphically illustrated in **Figure 5**) and asked which of them can better capture the firing rate variability in the biophysical models.

Specifically, four types of feedforward, backpropagation Artificial Neural Networks (ANNs) were constructed (see **Online Methods**). In the *2-layer modular ANN*, supralinear and sublinear dendrites were simulated as 2 parallel hidden layers consisting of a logistic sigmoid and a sublinear activation function *y(x) = (x+2)^0.7^-2*, respectively^49^ (**Figure 5**). The number of activated synapses allocated to supralinear vs. sublinear dendrites in the biophysical models was used as input to the respective hidden layers. The output layer represented the soma/axon of the biophysical model and consisted of a linear activation function. In the *linear ANN*, there was only a single hidden layer receiving input from all dendrites and consisting of linear activation functions (**Figure 5**). We also constructed two ANNs with the exact same architecture as the linear one, but with either a) a logistic sigmoidal (*2-layer supralinear ANN*) or b) a sublinear *y(x) = (x+2)^0.7^-2* (*2-layer sublinear ANN*) activation function in the hidden layer neurons (**Supplementary Figure 10**). These ANNs represent FS BCs with just one type of non-linear dendrites.

For all 8 FS BC model neurons the *linear* and *2-layer modular ANNs* were trained using the number of synapses to supra-/sublinear dendrites as inputs to the respective hidden layers and the mean firing rate of the soma as target output. A randomly selected 80% of our synaptic activation data set was used to train the model and the rest 20% to test its generalization performance (see **Online Methods**). Performance accuracy was estimated based on regression analysis between the ANN-generated firing rates and those produced by the biophysical models. Fits for two representative model cells are shown in **Figure 6a-d**, while the overall performance for all 8 model cells is shown in **Figure 6e. Figure 6f** demonstrates the performance of both ANN types for a dataset of the same power (number of inputs = 60), whereby the location of the inputs varies. As evident from the results, the *2-layer modular ANN* outperformed the *linear ANN* in all cases tested.

However, the performance of the *linear ANN* was relatively good. This can be attributed to the wide range of activated synapses (2 to 60) which resulted in large differences in the somatic firing, irrespectively of synapse location, and can thus be captured by any linear model (also see the work of Poirazi et al 2003b)^40^. Therefore, we also assessed the performance accuracy of *linear* and *2-layer modular ANNs* to the more challenging task of discriminating between input distributions corresponding to the exact same number of synapses. To do so, we subdivided the data into input categories corresponding to 20, 40 and 60 synapses, respectively. In these more challenging conditions, the *2-layer modular ANN* clearly outperformed the respective *linear ANN*, which failed to explain the variance produced by differences in input location. This result was consistent for all model cells as shown in **Table 1**. Performance for the 60-synapse case is shown in **figure 6f**.

Taken together, this analysis suggests that a 2-layer artificial neural network that considers both types of dendritic non-linearities is a much better mathematical abstraction for FS basket cells than the currently assumed linear point neuron.

### Bimodal nonlinear integration of FS basket cells enhances memory encoding

In order to investigate the functional implications of our findings, we built a canonical microcircuit network model^41^ composed of simplified 2-stage excitatory neurons, FS BCs and dendrite targeting (SOM+) interneurons (**Figures 1d and Supplementary Table 6)**. The model includes inhibitory feedback connectivity, multi-dendrite and perisomatic interneurons. It implements plasticity-related processes which act on multiple temporal and spatial scales: two-stage dendritic integration, dendritic calcium dynamics, synaptic tagging and capture (STC), CREB-dependent excitability and homeostasis (see **Online Methods**).

The network model was first trained to encode a single memory^41^ (see **Online Methods**) using FS BCs with either a) purely linear or b) bi-modal (sublinear and supralinear) dendritic subunits, as predicted by the compartmental modeling analysis (**Figure 2**). SOM+ interneurons were modeled as having either sublinear, linear, supralinear, or bi-modal dendritic subunits (**Supplementary Figure 11**). In these simulations, synaptic inputs to the FS BCs cells were either a) randomly distributed in all dendrites (Dispersed) or b) clustered within 33% of all dendrites (Clustered) (see **Online Methods**). The properties of the resulting memory engram (*i.e*. the population of active excitatory neurons during recall) were assessed by analyzing the activity of excitatory neurons during recall 24 hours after the learning event (**Figure 7e**).

Our results indicate that, compared to linear dendrites, bi-modal FS BCs dendrites lead to significant reductions in the size of the resulting memory engram (p-value=5.8e-15), and the mean engram firing rates (p-value=3.1e-18 linear-dispersed, p-value=7.2e-10 linear-clustered) (**Figure 7a,b**) while they also increase the network firing sparsity (p-value=0.00095 linear-disperse, p-value =0.00338 linear-clustered) (**Figure 7c**). All of the above suggest that dendritic bi-modality in FS BCs promotes resource savings in the encoding of new memories.

As predicted by our multi-compartmental models (**Figure 4**), we found that dispersed synaptic activation is beneficial to engram properties by further reducing the engram size (p-value=4.0e-6) and the mean firing rate (p-value=1.7e-7) (**Figure 7a-b**). Summarizing, the memory engram properties indicate that bi-modal FS BC dendrites receiving dispersed inputs confer resource consumption advantages to memory encoding by a) increasing the sparsity of the population, b) recruiting fewer engram neurons and c) reducing the overall network excitability. The above findings were unaffected by the presence of either linear, supralinear or bi-modal SOM+ model dendrites (**Supplementary Figure 11**).

Finally, we also assessed the role of FS-BC nonlinearities in memory linking, by encoding two memories separated by 1 or 24 hours in the same network model and measuring the population overlap of the resulting memory engrams. According to previous work^33,41,50^, memories learned in close temporal proximity (e.g. 1 hour apart) display increased engram overlap compared to distant memories (24 hours apart). Overlapping storage is also associated with behavioral binding of the two memories and has been proposed to underlie the linking of memories across time^51^. We found that linear FS BC dendrites result in substantially larger engram overlaps in the circuit model compared to bi-modal dendrites (**Figure 7d**), for the 1-hour case. These overlaps are in fact significantly larger than the experimentally reported ones^50^ (~20%), suggesting that the two memories may interfere with one another. Taken together, our network modeling analysis suggests a beneficial role of nonlinear dendrites in FS BCs with respect to memory encoding, storage capacity as well as the binding of memories over time.

## DISCUSSION

The role of dendrites in interneuron computations is a rapidly emerging and debatable subject^52^. Several recent reports present exciting findings according to which dendrites may serve as key players^2,11,37,38,53^. For example, sodium spikes and supralinear calcium accumulation have recently been reported in the dendrites of FS BCs^2^, yet the consensus still favors the linear point neuron dogma^6,45,52^. The present study provides new insight into this ongoing debate by systematically analyzing the dendritic integration mode of FS BCs in two brain areas: The Hippocampus and the PFC. We do so using an extensive set of computational tools that extends from detailed biophysical single cell models, to reduced integrate-and-fire single cell and circuit models as well as artificial neural network models (**Figure 1**). We predict that dendrites of both cortical and hippocampal FS BCs operate in one of two modes of synaptic integration: supralinear or sublinear (**Figure 2**). Supralinearity is due to the generation of dendritic sodium spikes (**Supplementary Figures 7)**, which are in turn gated by the morphology (**Figure 3**) of dendrites. Moreover, we find that somatic output is influenced by the spatial distribution of activated synapses, with dispersed input inducing higher firing rates than clustered activation. This feature is opposite to pyramidal neurons^40^ and is attributed to a) the presence of sublinear dendrites in FS BCs and b) the small dendritic diameter, increased A-type current and fast EPSP kinetics of cp-AMPA receptors found in these cells^6^ (**Figure 4, Supplementary Figure 12)**. Due to these properties, a 2-layer modular Artificial Neural Network abstraction with both sub- and supra-linear hidden neurons (**Figure 5**) captures the spiking profile of biophysical neurons with much higher accuracy than a linear ANN, analogous to a point neuron. This is true for all of the 8 morphological reconstructions of FS BCs tested and is more evident for datasets in which the number of inputs is fixed but their location varies (**Figure 6, Table 1**). This is because discriminating the effect of input location as opposed to input strength is a much more challenging task and pushed the linear ANN to its performance limits. Finally, we show that such a 2-stage integration model facilitates the efficient encoding, storage and discriminability of memories in a biologically relevant circuit model across time (**Figure 7**).

### Mediators of supralinear and sublinear dendritic integration in FS basket cells

A bimodal dendritic integration is predicted for all hippocampal and PFC morphologies analyzed. Supralinearity was found to be due to the occurrence of dendritic sodium spikes (**Supplementary Figure 7**). Several mechanisms can influence the generation of such dendritic spikes: ionic conductances (primarily of sodium currents but also potassium currents) and morphological features. In our models, biophysical mechanisms are constrained by existing experimental data and dendritic sodium conductances are kept to a minimum (10 times smaller than the soma^6^), so as to minimize the probability of non-physiological dendritic spiking. Sensitivity analysis further demonstrates that results are robust to physiological variations in a wide range of active dendritic conductances (**Supplementary Figure 6**). These findings strongly suggest that dendritic spiking in certain dendrites of FS basket cells are highly likely to occur under physiological conditions, in line with recent experimental reports^2^.

Apart from sodium currents as a universal enabling mechanism, we find a key role of morphology in gating local dendritic spikes. A combination of dendritic length and mean diameter, or otherwise the dendritic volume and input resistance, is statistically different between sub-(smaller) and supralinear (larger) dendrites across all morphologies tested (**Figure 3**). The inability of small-volume dendrites (**Figure 3F**) to support sodium spikes is attributed to their high input resistance (**Figure 3F**), fast kinetics of calcium permeable AMPA receptors and the high density of A-type potassium channels^6,12^. This combination results in large, fast EPSPs that are very efficient in activating IA currents, which in turn repolarize the membrane^6,12^. This mechanism has been previously proposed by others^6,12,42^, is supported by our morphology and IA manipulation experiments (**Figure 4**) and is in line with other studies reporting a similar effect of morphology on the ability of dendrites to generate local spikes^36^.

### Functional coexistence of sub- and supra-linear dendrites within FS basket cells

Our simulations predict the co-existence of both sublinear and supralinear dendrites in all FS BCs models (**Figure 2, Supplementary Figures 4–9**). Similar bimodal dendritic integration has been reported in hippocampal CA1 pyramidal neurons^35,44^ and predicted in PFC pyramidal neurons^48^.

The existence of sublinear dendritic branches supports the idea of inhibitory neurons acting as coincidence detectors by aggregating spatially disperse and nearly synchronous synaptic inputs^6^. Moreover, sublinear dendrites can compute complex non-linear functions similar to those computed by sigmoidal dendrites^49^, thus substantially extending the processing capacity of these neurons compared to a linear integrator. Why have two types of nonlinearity then?

Our network modeling predicts that the presence of both types of nonlinearities confer substantial benefits to network computations and especially to memory encoding. We find that bimodal nonlinearities in the dendrites of FS BCs, enables the encoding of new memories within a smaller neuronal population, thus increasing sparsity and storage capacity. These nonlinearities also facilitate the interaction of memories over time, via decreasing the possibility of interference (**Figure 7**).

### Not that Simple: FS basket cells as 2-layer modular ANNs

Artificial Neural Network analysis demonstrates that a FS basket cell is better described by a 2-stage abstraction that incorporates both modes of dendritic integration (**Figure 5,6**). This work, along the lines of the 2-stage model proposed for pyramidal neurons^40^, strongly challenges the prevailing point neuron dogma. The 2-stage abstraction is supported by experimental reports of dendritic sodium spikes and supralinear calcium accumulations^2^ while it also explains sublinear dendritic integration^6,11,29,55^, providing a unifying framework for interneuron processing.

Possible limitations of our work include the imprecise modeling of ionic and synaptic mechanisms given the shortage of sufficient information for FS BCs models. This limitation is counteracted by the sensitivity analysis of the mechanisms that mostly influence our findings and their consistency across several cortical and hippocampal morphologies. Another limitation is the lack of inhibitory inputs (except from the autaptic GABAa current that is incorporated in all models). Inhibitory inputs consist of just 6% of all incoming contacts in Fast Spiking interneurons^6,56,57^. Thus, our results are unlikely to be affected by inhibitory inputs. FS basket cells in the hippocampus and the neocortex are highly interconnected by gap junctions^6^, that can speed the EPSP time course, boost the efficacy of distal inputs and increase the average action potential frequency after repetitive synaptic activation^6^. All of these effects would contribute to stronger responses but unless gap junctions are spatially specific to certain branches and not others, they are unlikely to influence the non-linear integration modes of dendrites.

## Conclusion

This work provides a novel view of dendritic integration in FS basket cells, that extends in hippocampal and cortical areas^41^. Here we suggest new reductionist models for interneuron processing, in which dendrites play a crucial role. Experimental validation of these new models is likely to change the way we think about interneuron processing, attribute new and exciting roles to FS basket cells and open new avenues for understanding interneuron contributions to brain function.

## Acknowledgments

This work was supported by the ERC Starting Grant dEMORY to Panayiota Poirazi. Alexandra Tzilivaki was supported by the Google EMEA Scholarship, by the Onassis Foundation Scholarship and by the Einstein Foundation Berlin Fellowship. The authors would like to thank Stefanos Stamatiadis for providing feedback on simulation procedures and all the members of the Computational Biology Lab for helpful discussions.

## AUTHOR CONTRIBUTIONS

AT and PP designed the experiments. AT and GK performed the simulations and analyzed the data. AT and PP wrote the manuscript. All authors edited the manuscript. PP supervised the work.

## Supplementary Figures

**Supplementary Figure 1.**
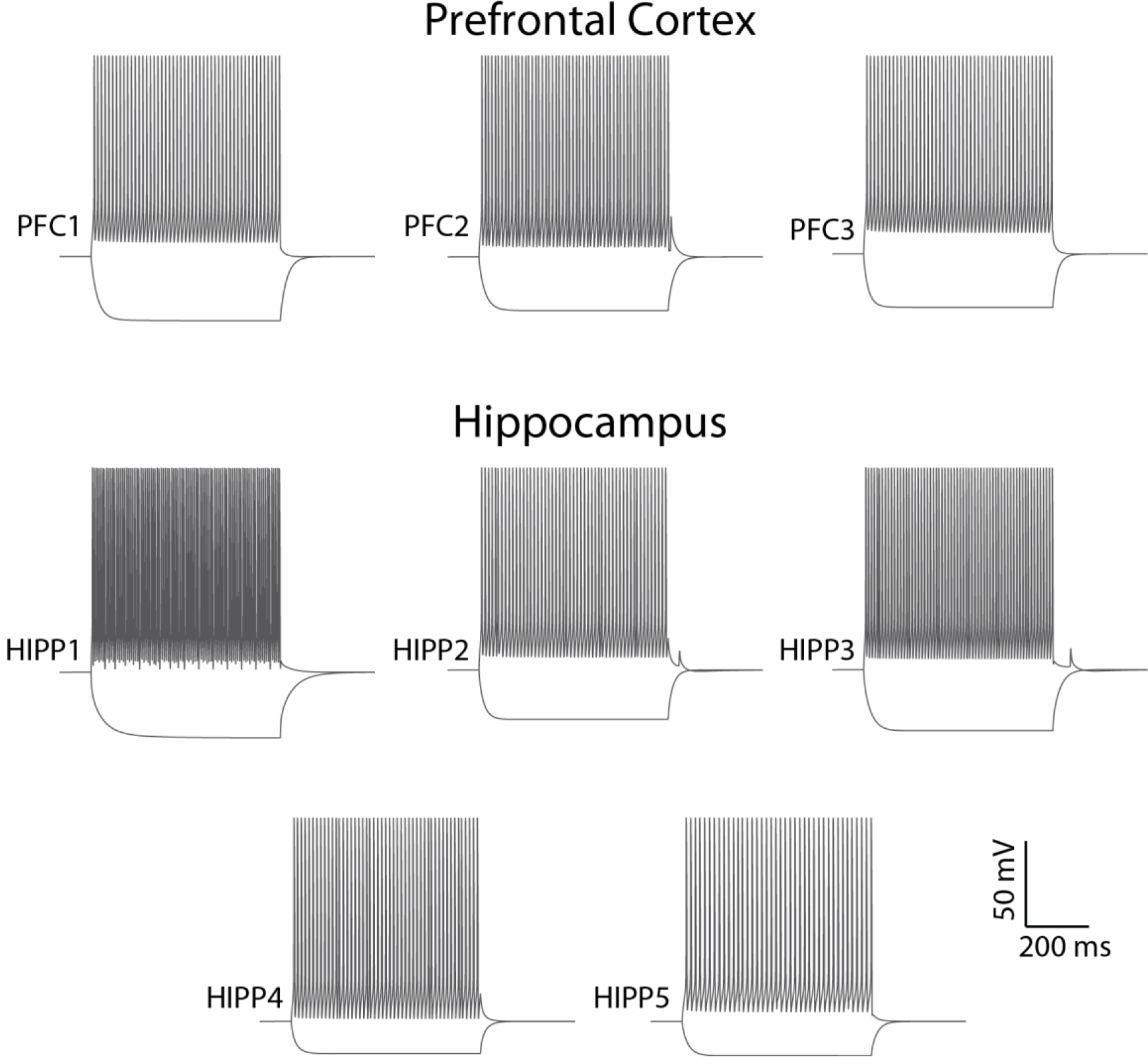
Model cell firing profiles. Somatic Current-clamp traces of Hippocampal (A) and PFC (B) model cells, after a depolarizing current injection in somata (500 pA; 1000 ms) evoked a high-frequency firing pattern. A hyperpolarizing current injection in somata (−300pA, 1000ms) induced a realistic hyperpolarizing response.

**Supplementary Figure 2.**
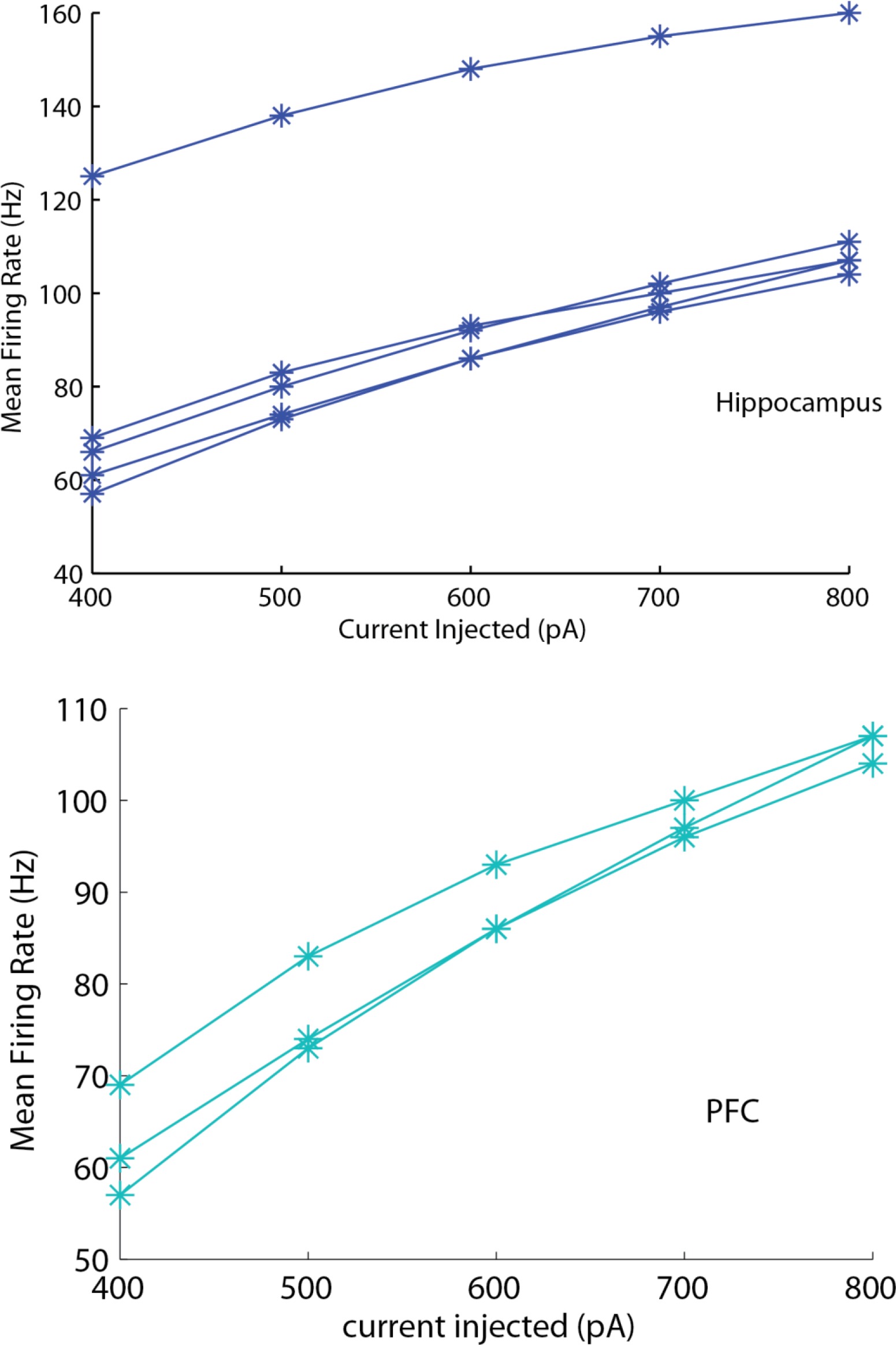
Mean firing frequencies in response to injected currents of different amplitudes (600 ms duration) in Hippocampal (up) and PFC (down) model cells.

**Supplementary Figure 3.**
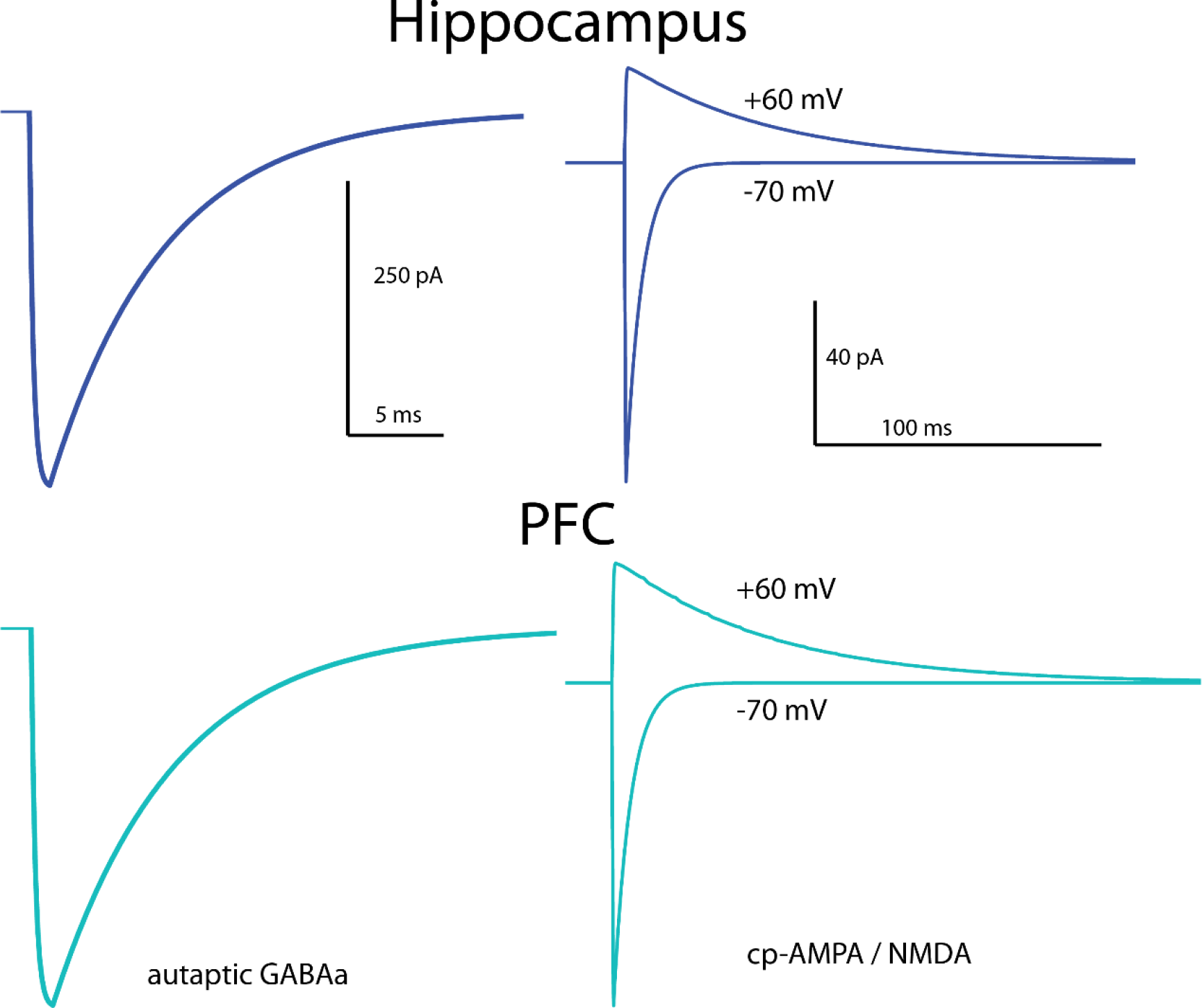
Validation of synaptic currents in Fast Spiking basket cells. Left. A three-step voltage clamp of voltage changes from −70 mV to 10 mV (duration 1 ms) and back to −70 mV was used to produce a self-inhibitory (autaptic) current. During the validation of this current, the reversal potential of Cl− was adjusted from −80 to −16 mV, in order to reproduce the experimental set up of Bacci et al., 2003. However, a physiological reverse potential (−80 mV) was used for all other simulations. Right. Model reproduction of cp-AMPA (−70 mV) and NMDA (+60 mV) currents in response to stimulation of 2 synapses as per Wang et al., 2009. * each trace represents the mean of all Hippocampal and PFC cells respectively.

**Supplementary Figure 4.**
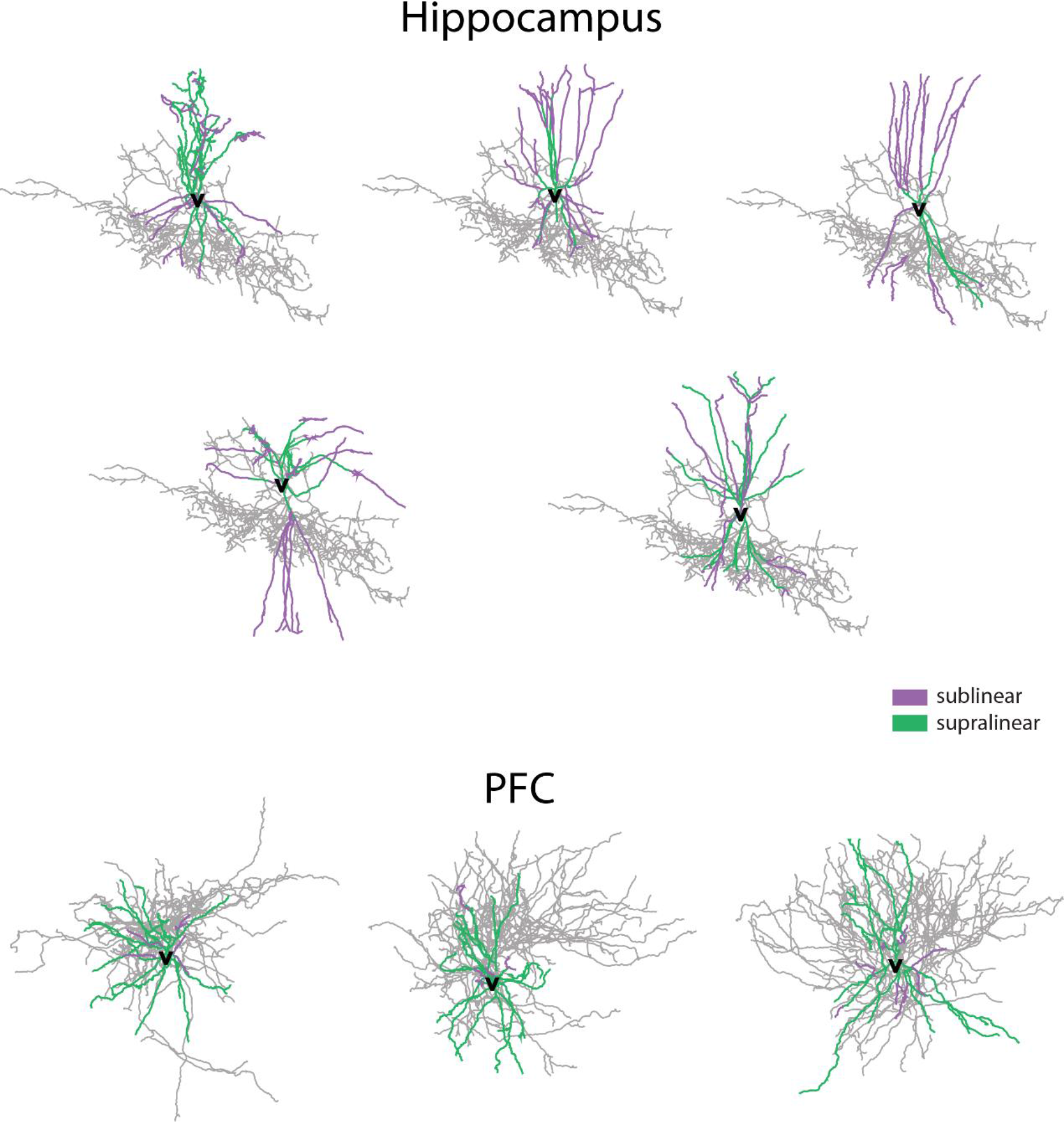
Related to **Figure 2**. Bimodal non-linear integration in Fast Spiking basket cells. Supralinear (blue) and sublinear (magenta) dendrites shown in each model cell.

**Supplementary Figure 5.**
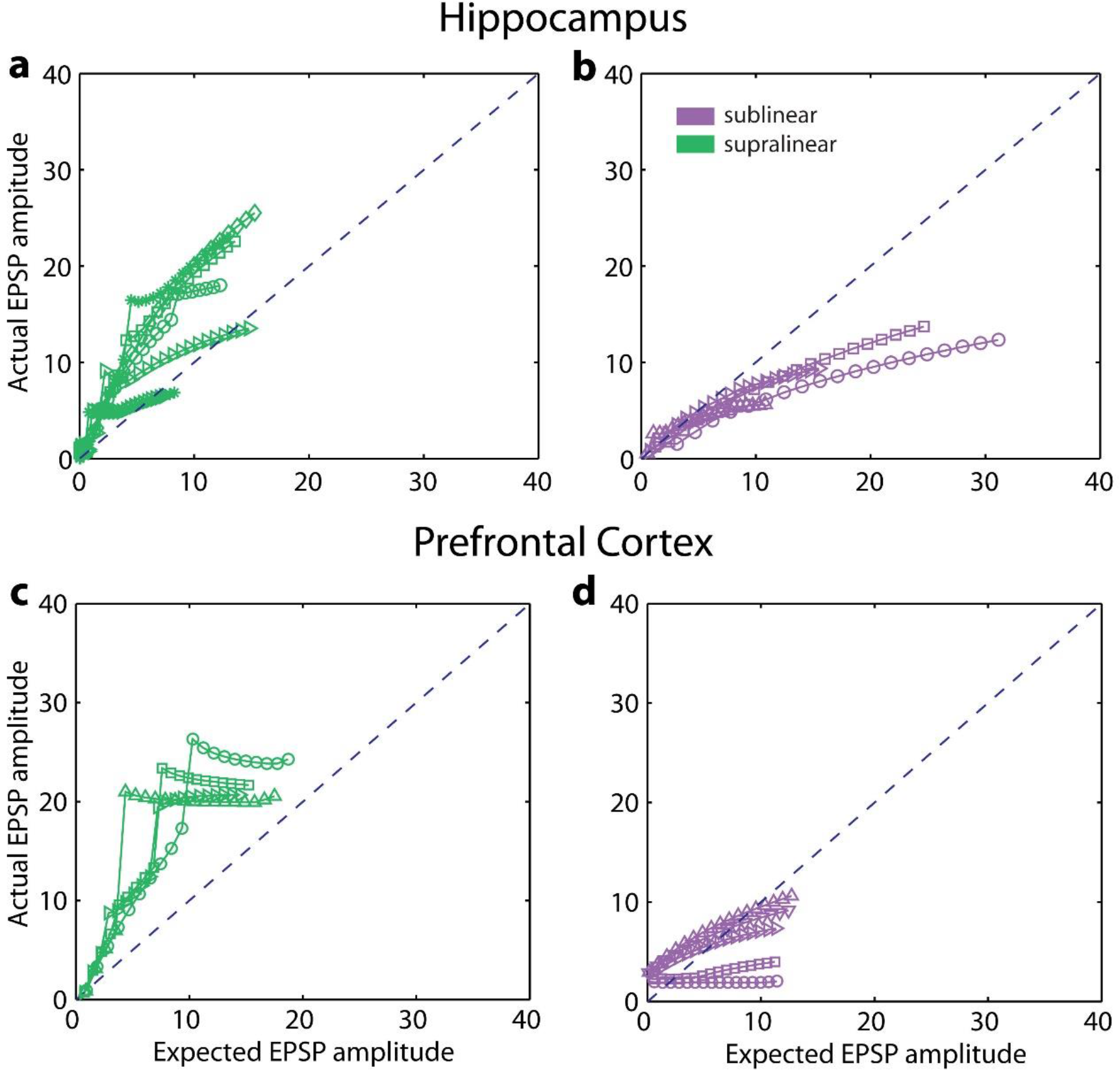
Related to **Figure 2**. Bimodal non-linear integration in Fast Spiking basket cells. Representative Somatic EPSPs after stimulation (single pulse) of an increasing number of synapses (1:1:20), uniformly distributed within dendrites.

**Supplementary Figure 6.**
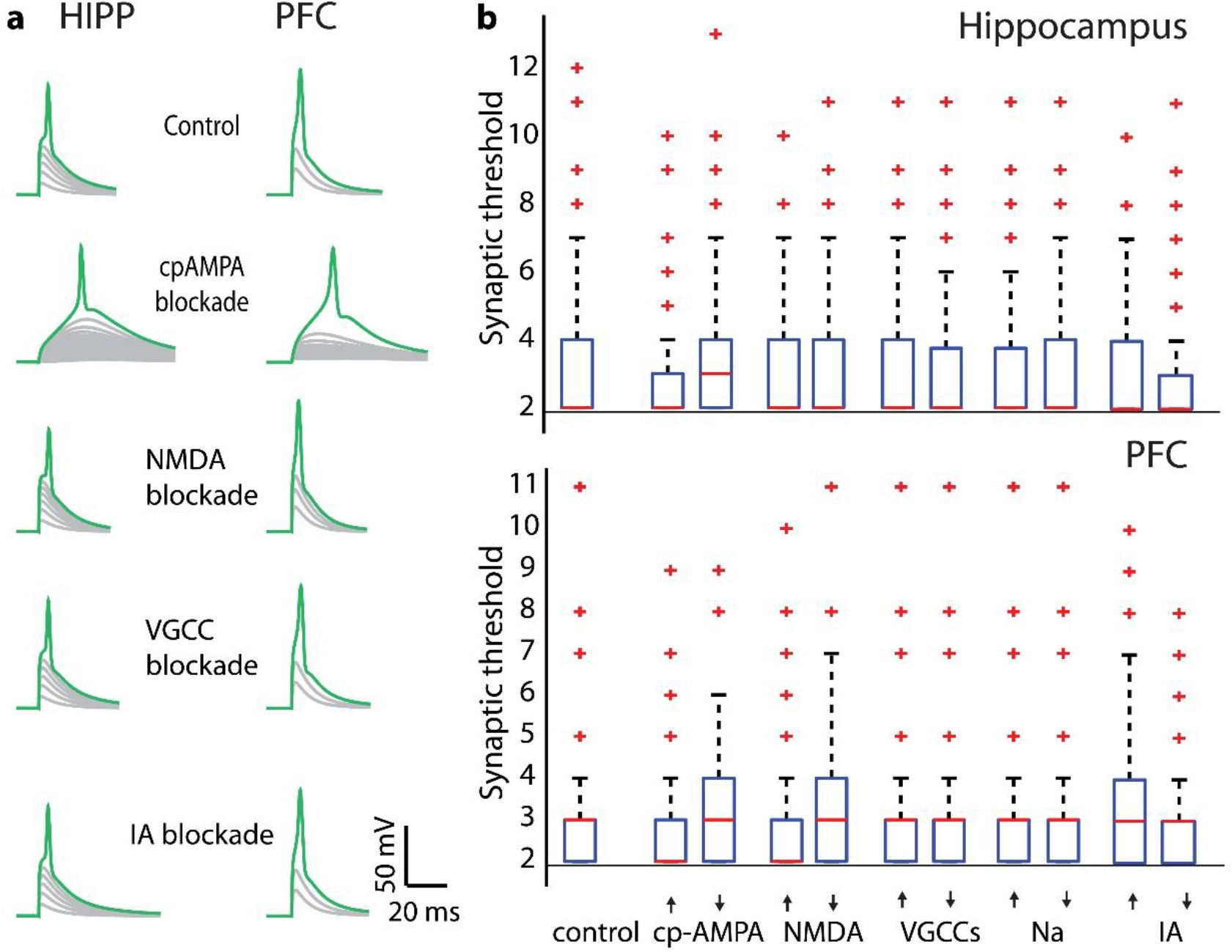
Related to Figure 1. **a.** Presence of supralinear summation in dendrites of FS BCs after blockade of multiple active currents respectively in Hippocampus (left) and PFC (right). **b.** Sensitivity analysis of biophysical dendritic mechanisms reveals minor changes in the synaptic threshold for spike generation in supralinear dendrites across Hippocampus and PFC.

**Supplementary Figure 7.**
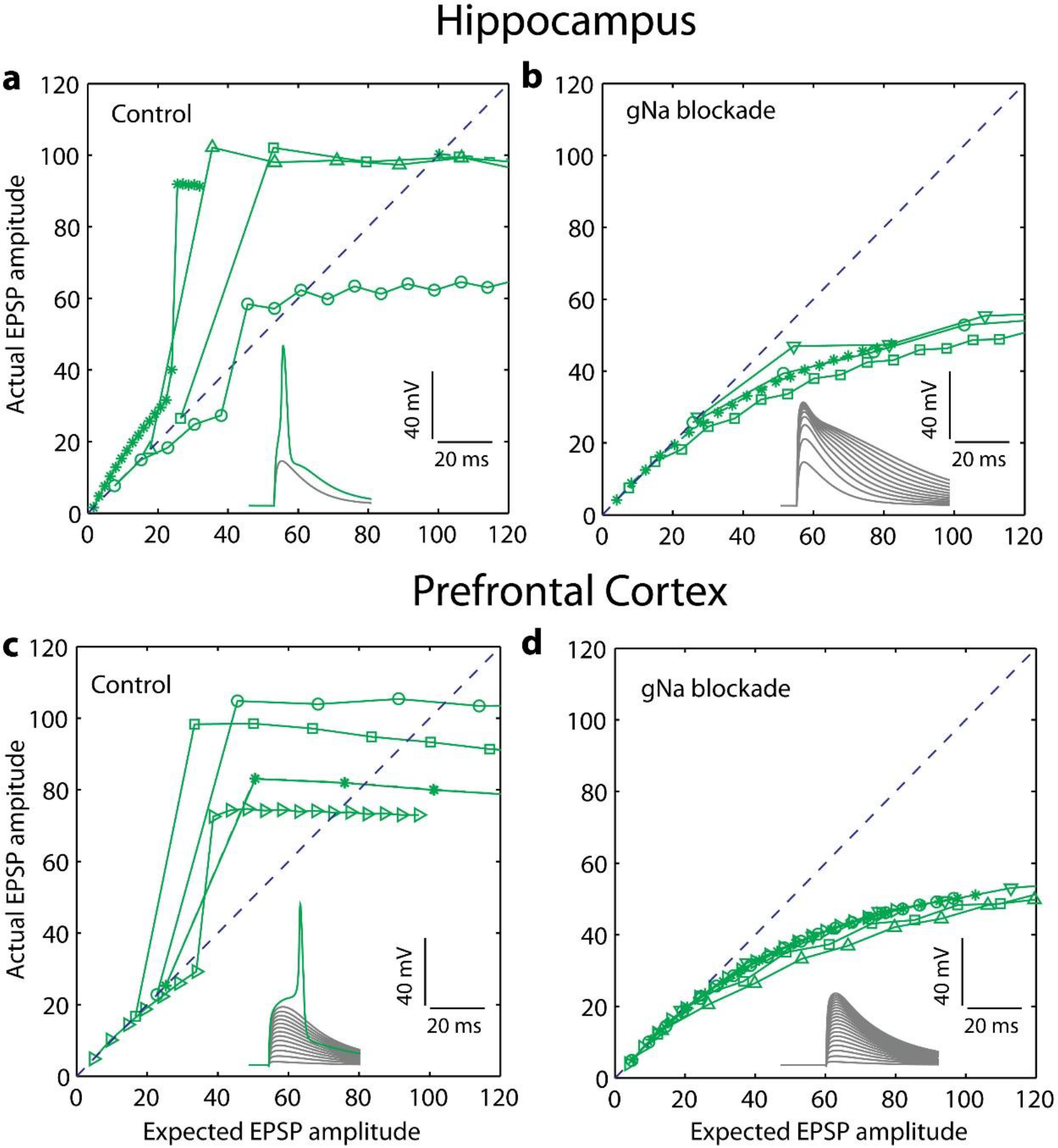
Related to Figure 1. Blockade of active sodium conductances in the dendrites of FS BCs, totally eliminates the supralinear operation mode. Hippocampal (**a**) and PFC (**c**) representative supralinear responses of dendrites under physiological conditions. Dendritic spikes are eliminated both in Hippocampal (**b**) and PFC (**d**) FS BCs dendrites EPSP responses after blockade of sodium conductances. Linear line represents linear summation.

**Supplementary Figure 8.**
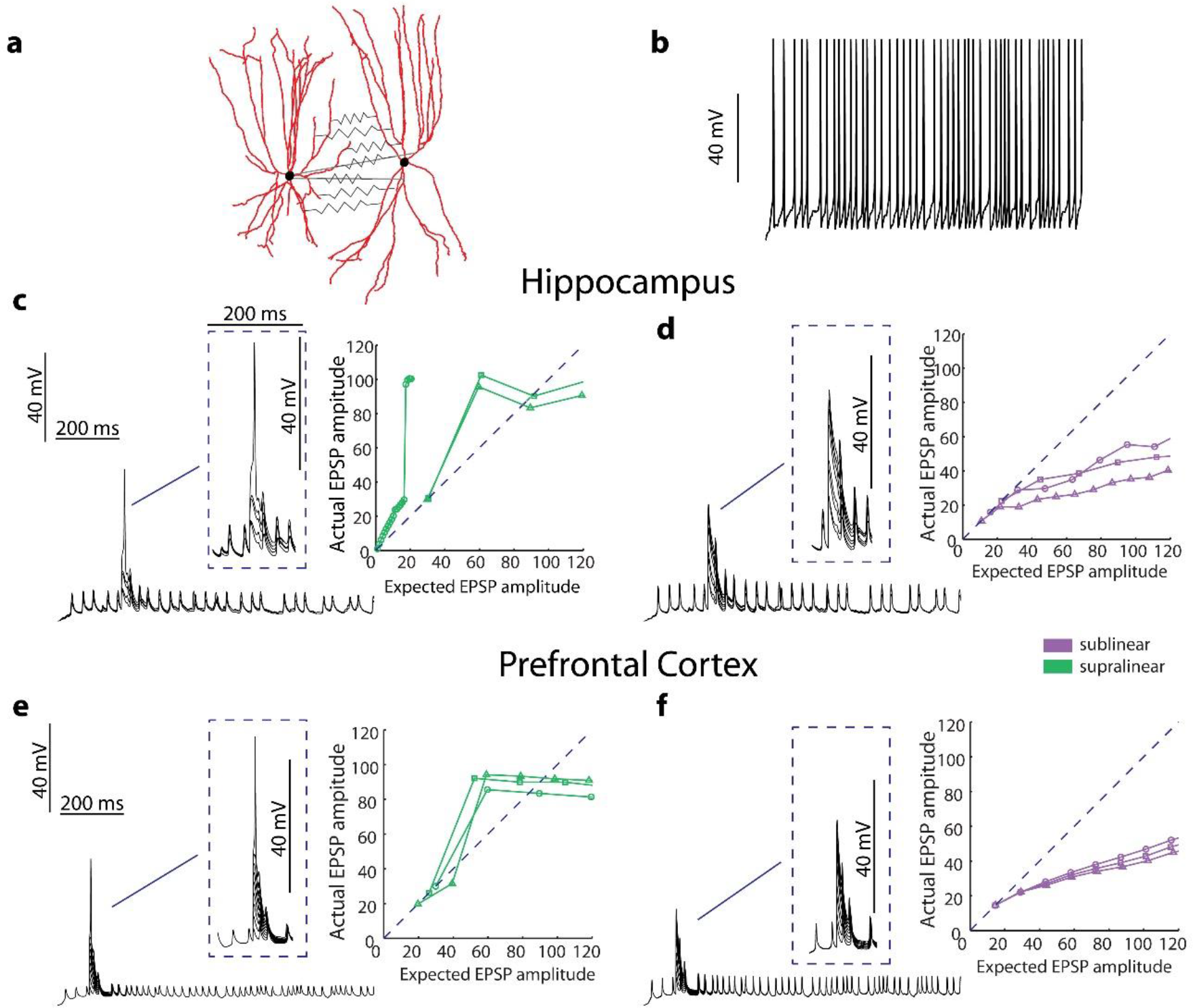
FS BCs exhibit supralinear and sublinear dendritic responses in the presence of Gap Junctions. A) Illustrated dendritic trees that are interconnected with Gap Junctions. B) Presynaptic firing rate (~30 Hz). Supralinear (C,E) and sublinear (D,F) dendrites co-exist in Hippocampal (up) and PFC (down) Fast spiking basket cells models.

**Supplementary Figure 9.**
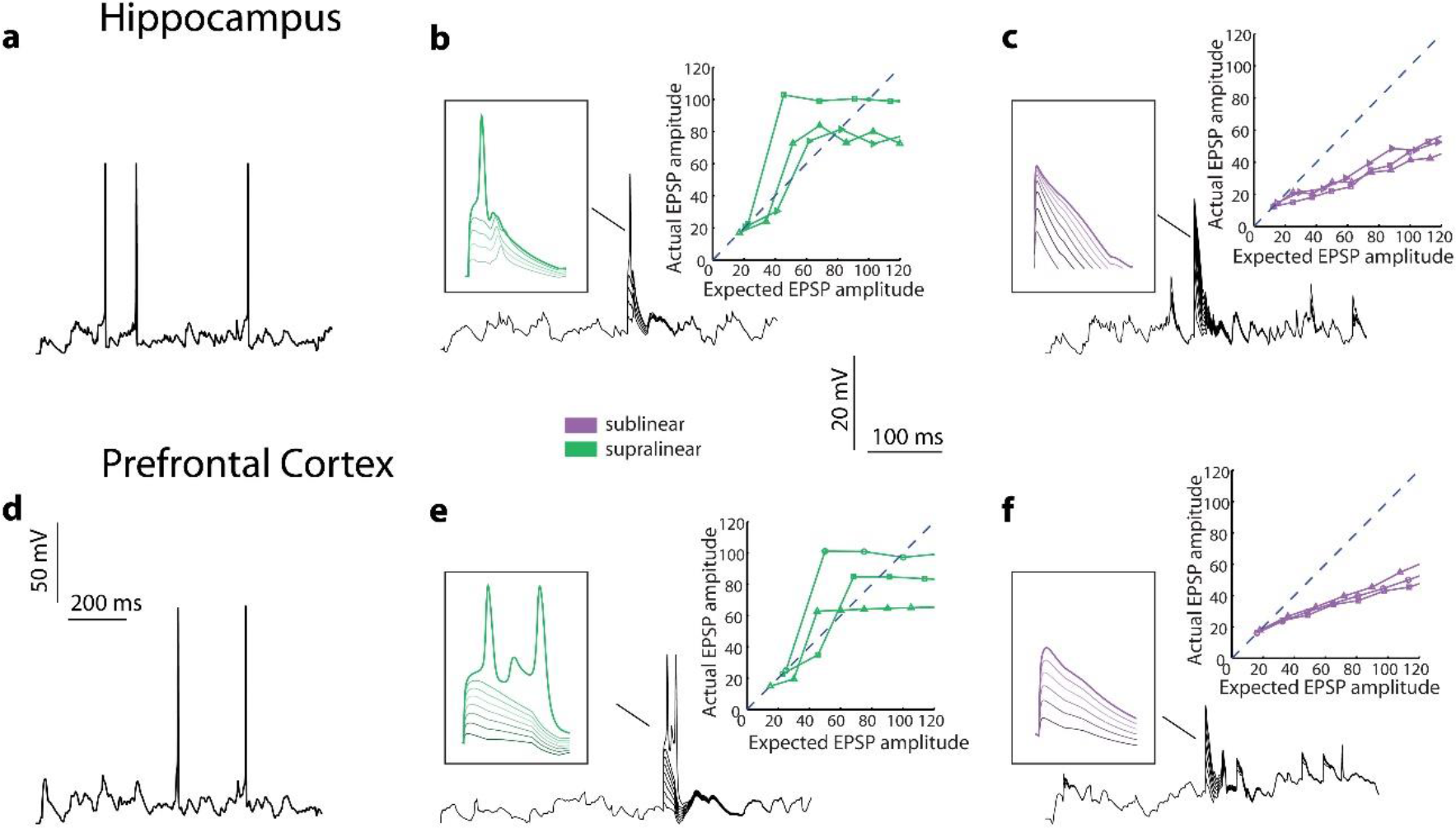
FS BCs exhibit supralinear and sublinear dendritic responses in the presence of *in vivo*-like fluctuations. Somatic firing rate of 3±1 Hz induced in Hippocampal (A) and PFC (D) models of FS BCs after synaptic activation of randomly selected dendrites with 10Hz Poisson spike trains. Supralinear (B,E) and sublinear (C,F) dendrites co-exist in FS BCs of Hippocampus (up) and PFC (down).

**Supplementary Figure 10.**
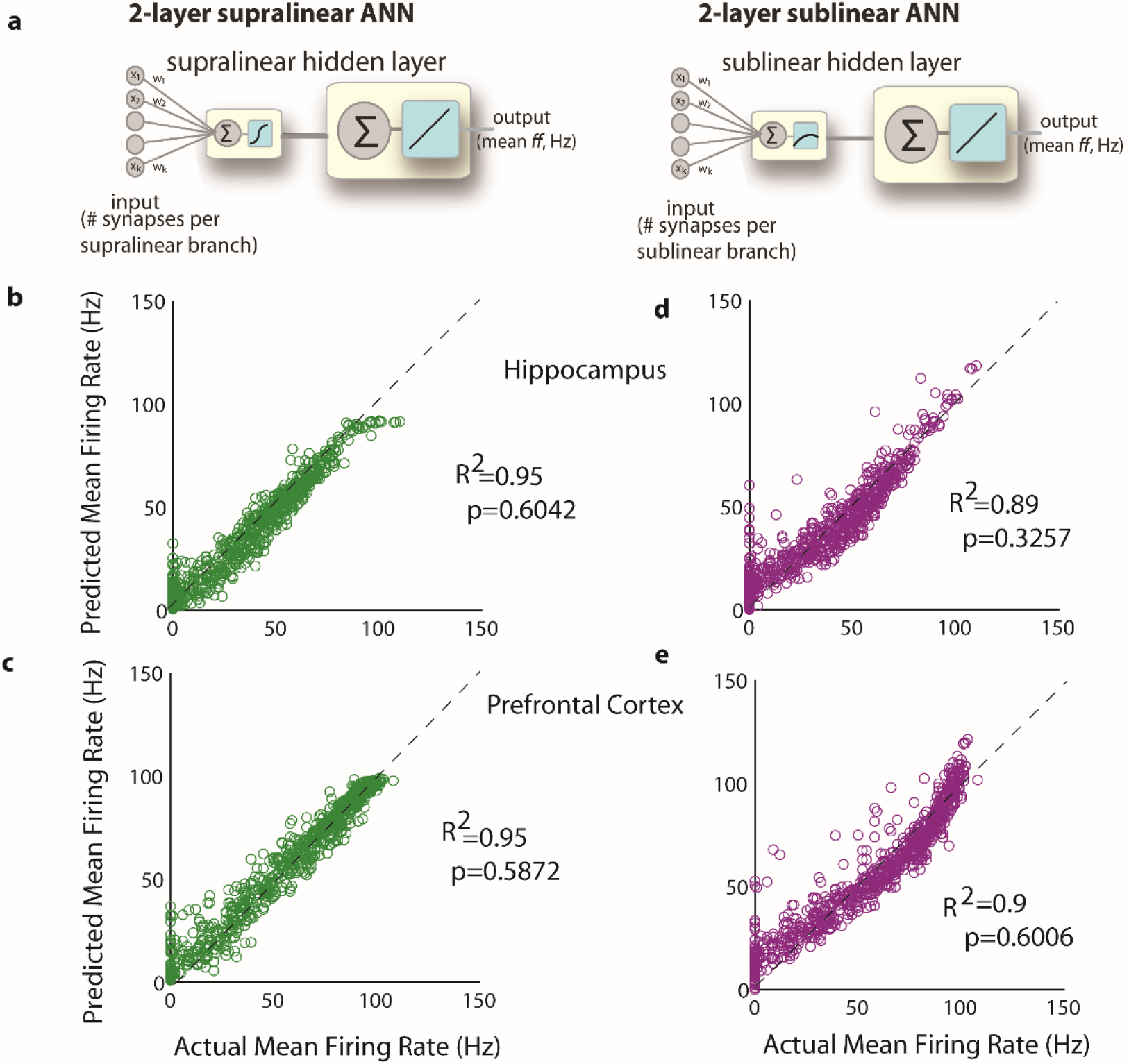
Related to **Figure 6**. Linear regression analysis for one hidden layer supralinear (**b**,**c**) and one hidden layer sublinear (**d**,**e**) ANNs for one indicative Hippocampal (top) and one indicative PFC (bottom) model cell. Actual Mean Firing Rates (Hz) correspond to the responses of the compartmental model when stimulating -with 50Hz Poisson spike trains-varying numbers of synapses (1 to 60), distributed in several ways (clustered or dispersed) within both sub- and supra-linear dendrites. Expected Mean Firing Rates (Hz) are those produced by the respective ANN abstraction when receiving the same input (number of stimulated synapses) in its respective sub-/supra- or linear input layer nodes.

**Supplementary Figure 11.**
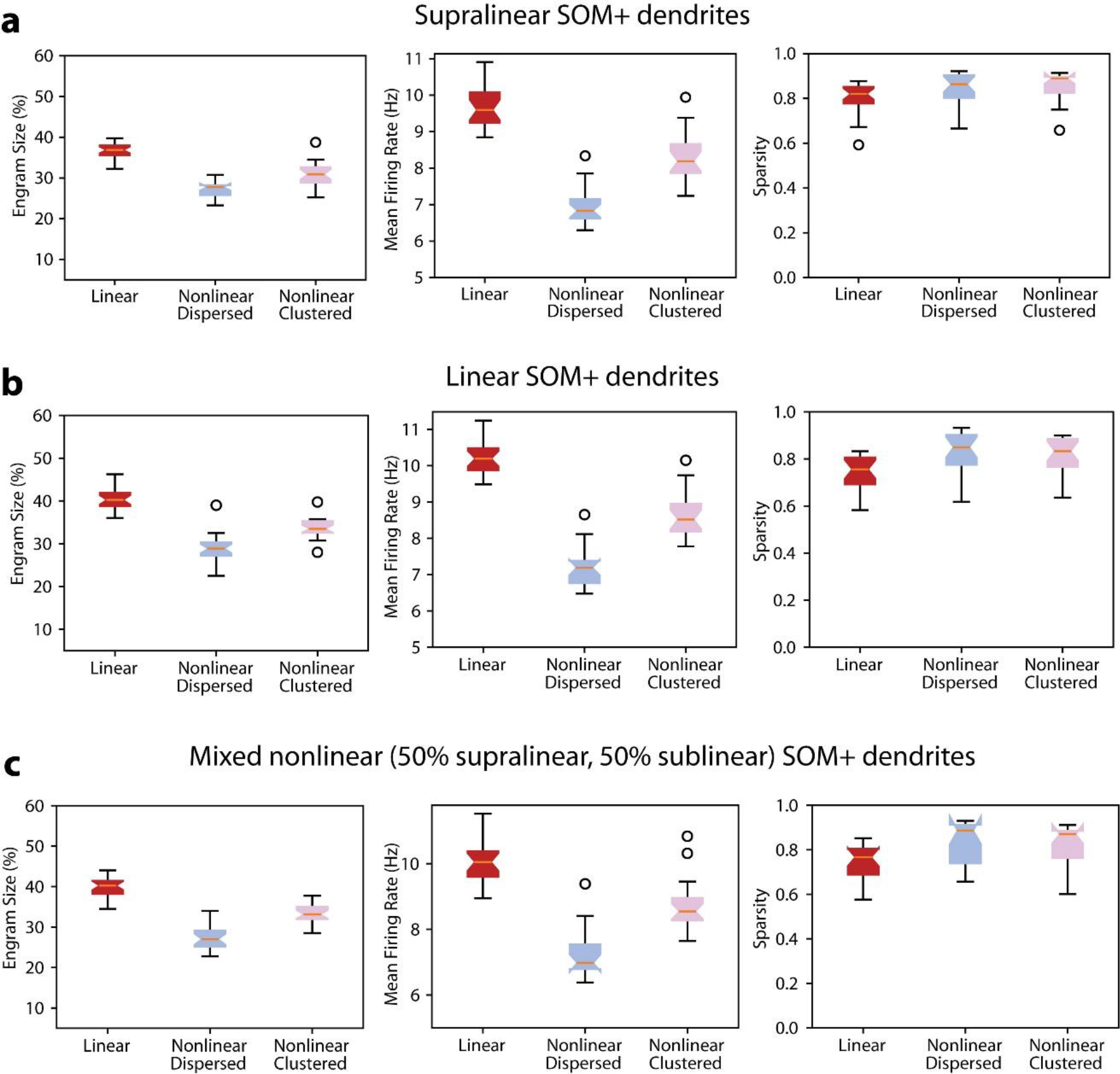
Related to **Figure 7.** Manipulation of SOM+ models dendritic transfer function results in almost identical responses of multiple properties of the canonical microcircuit. Modeled SOM+ dendrites A) Supralinear B) Linear C) Supralinear and sublinear (50% of each mode)

**Supplementary Figure 12.**
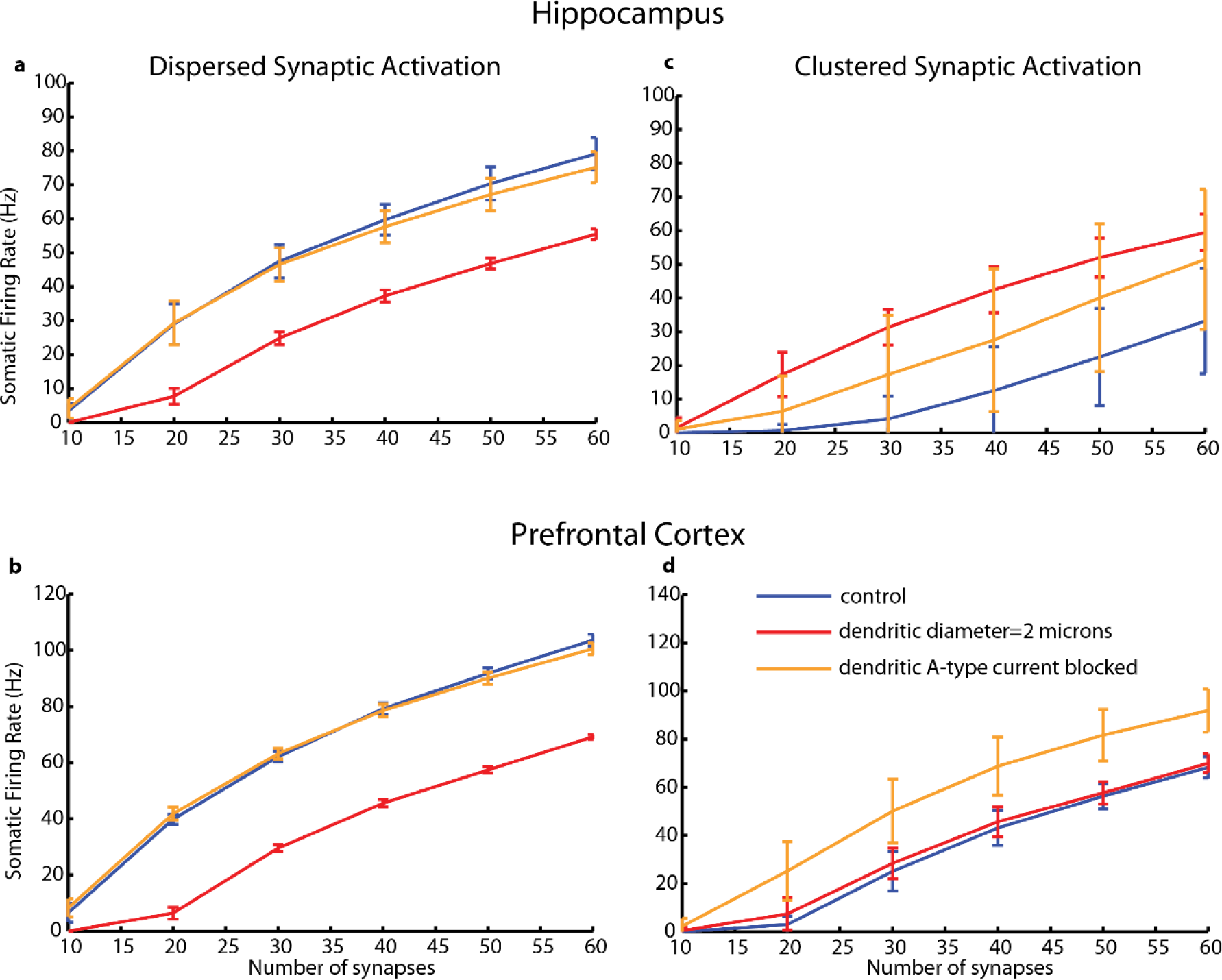
Related to **Figure 4:** Firing rate responses (in Hz) from one Hippocampal (**a**, **c**) and one PFC (**b**, **d**) model cell, in response to stimulation of increasing numbers of synapses (10 to 60) that are either randomly distributed throughout the entire dendritic tree or clustered within a few dendritic branches. Effect of dendritic diameter (red, setting the diameter of all dendrites to 2 microns) and A-type current (orange, setting the conductance of dendritic A-type currents to zero) on somatic firing rates in response to synaptic stimulation under dispersed and clustered spatial arrangements. As shown in panels **a**, **c** disperse synaptic arrangements benefit mostly from the dendritic morphology of FS BCs, as setting the diameter to 2 microns sharply decreases this preference. Clustered arrangements on the other hand (panels **b**, **d**) are severely hampered by the high conductance of the A-type potassium channels in these cells, as blockade of these currents enhances somatic output. This potassium current does not penalize disperse inputs as much, simply because it is not as strongly activated as in the case of clustered activation (which induces much higher local depolarizations and thus stronger A-type channel activation). (p-values for the various comparisons: hippocampus, disperse: diameter vs. control =0.0018, IA vs. control, non-significant; hippocampus, clustered: diameter vs. control=0.0048, IA vs. control=0.0087; PFC, disperse diameter vs. control =0.0014, IA vs. control, non-significant; PFC, clustered diameter vs. control =0.0102, IA vs. control=0.0026)

